# GFAT2 and AMDHD2 act in tandem to control the hexosamine biosynthetic pathway

**DOI:** 10.1101/2021.04.23.441115

**Authors:** Virginia Kroef, Sabine Ruegenberg, Moritz Horn, Kira Allmeroth, Lena Ebert, Seyma Bozkus, Stephan Miethe, Bernhard Schermer, Ulrich Baumann, Martin S. Denzel

## Abstract

The hexosamine biosynthetic pathway (HBP) produces the essential metabolite UDP-GlcNAc and plays a key role in metabolism, cancer, and aging. The HBP is controlled by its rate-limiting enzyme glutamine fructose-6-phosphate amidotransferase (GFAT) that is directly inhibited by UDP-GlcNAc in a feedback loop. HBP regulation by GFAT is well studied but other HBP regulators have remained obscure. Elevated UDP-GlcNAc levels counteract the glycosylation toxin tunicamycin (TM) and thus we screened for TM resistance in haploid mouse embryonic stem cells (mESCs) using random chemical mutagenesis to pinpoint new HBP regulators. We identified the N-acetylglucosamine deacetylase AMDHD2 that catalyzes a reverse reaction in the HBP and its loss strongly elevated UDP-GlcNAc. To better understand AMDHD2, we solved the crystal structure and found that loss-of-function is caused by protein destabilization or interference with its catalytic activity. Finally, we show that mESCs express AMDHD2 together with GFAT2 instead of the more common paralog GFAT1. Compared with GFAT1, GFAT2 had a much lower sensitivity to UDP-GlcNAc inhibition, explaining how AMDHD2 loss-of-function resulted in HBP activation. This HBP configuration in which AMDHD2 serves to balance GFAT2 activity was also observed in other mESCs and, consistently, the GFAT2/GFAT1 ratio decreased with differentiation of mouse and human embryonic stem cells. Together, our data reveal a critical function of AMDHD2 in limiting UDP-GlcNAc production in cells that use GFAT2 for metabolite entry into the HBP.

## Introduction

The hexosamine biosynthetic pathway (HBP) is an anabolic branch of glycolysis consuming about 2-3% of cellular glucose^1, 2^. It provides substrates for various posttranslational modification reactions and has been strongly associated with stress resistance and longevity as well as cell growth and transformation^3–5^. Thus, the HBP plays an essential role for metabolic adaptations and cellular homeostasis^6^.

In the first and rate limiting step of the HBP, glutamine fructose-6-phosphate amidotransferase (GFAT) converts fructose-6-phosphate (Frc6P) and L-glutamine (L-Gln) to D-glucosamine-6-phosphate (GlcN6P). The two mammalian GFAT paralogs GFAT1 and GFAT2 show 75-80% amino acid sequence identity^7^. While GFAT1 is ubiquitously expressed, GFAT2 is reported to be predominantly expressed in the nervous system. In the second step of the HBP, glucosamine-phosphate N-acetyltransferase (GNA1) acetylates GlcN6P to N-acetylglucosamine-6-phosphate (GlcNAc-6P) using acetyl-CoA as the acetyl donor. After isomerization into GlcNAc-1-phosphate (GlcNAc-1P) mediated by GlcNAc phosphomutase (PGM3), UTP is used in a final step by UDP-N-acetylglucosamine pyrophosphorylase (UAP1) to synthesize the final product UDP-GlcNAc. The HBP is the only source for UDP-GlcNAc and relies on substrates from carbon, nitrogen, fatty-acid, and energy metabolism. It is therefore optimally positioned as a metabolic sensor that can modulate downstream cellular signaling through UDP-GlcNAc dependent posttranslational modifications (PTMs)^1^.

UDP-GlcNAc is a precursor of several important biomolecules such as chitin, peptidoglycans, glycosaminoglycans, and for a number of dynamic glycosylation events. Mucin-type O-glycosylation plays an important role in the extracellular matrix^8^. N-linked-glycosylation orchestrates protein folding in the endoplasmic reticulum (ER) and is therefore crucial in protein homeostasis^9^. N-glycans further contribute to the cell surface glycocalyx as structural components of proteins^10^. Finally, the addition of single GlcNAc moieties to Thr/Ser residues, termed O-GlcNAcylation, occurs dynamically on hundreds of proteins thus modulating a variety of downstream pathways^11^. Surprisingly, this dynamic PTM is accomplished by a single protein, O-GlcNAc transferase (OGT), and O-GlcNAcase (OGA) is the only known enzyme to remove O-GlcNAc modifications^12, 13^. While it is known that these glycosylation reactions are limited by intracellular UDP-GlcNAc, how the HBP is regulated to adapt UDP-GlcNAc levels according to nutrient availability is poorly understood.

In a previous chemical mutagenesis screen in *Caenorhabditis elegans* we isolated mutants resistant to the toxin tunicamycin (TM) as a proxy for enhanced protein quality control and found that TM resistant mutants were enriched for longevity^3^. TM is a competitive inhibitor of UDP-GlcNAc:dolichylphosphate GlcNAc-1-phosphotransferase (GPT), which catalyzes the first step of N-glycan synthesis utilizing UDP-GlcNAc^14^. TM disrupts N-glycosylation and leads to proteins misfolding and proteotoxic stress^9^. We found that single amino acid substitutions in GFAT1 result in gain-of-function due to loss of UDP-GlcNAc feedback inhibition, elevating cellular UDP-GlcNAc levels and thereby counteracting TM toxicity^15^. By introducing the same gain of function mutation in GFAT1 of mouse neuroblastoma Neuro2a (N2a) cells, we confirmed a conserved mechanism^16^, suggesting that screening for TM resistance might be a suitable unbiased means to analyze the HBP through genetic approaches in mammalian cells.

In this study we combined chemical mutagenesis with whole exome sequencing in haploid murine cells and identified the N-acetylglucosamine-6-phosphate deacetylase AMDHD2 (Amidohydrolase Domain Containing 2) as a novel regulator of the HBP. Through AMDHD2 deletion, we discovered a configuration of the HBP that uses GFAT2 as the key enzyme. Functionally, GFAT2 shows a lower sensitivity to UDP-GlcNAc feedback inhibition compared to GFAT1 therefore requiring AMDHD2 to balance HBP metabolic flux.

## Results

### Chemical mutagenesis screen for tunicamycin resistance in mESCs identifies AMDHD2

Elevated HBP activity and high UDP-GlcNAc concentrations suppress TM toxicity, making TM resistance a proxy for HBP activity in genetic screens. To investigate HBP regulation in mammalian cells we therefore performed an unbiased TM resistance screen. The mutagen N-ethyl-N-nitrosourea (ENU) induces single nucleotide variants that enable a screen at amino acid resolution. Thus, we used ENU in haploid cells, which uniquely enable identification of recessive alleles^17–19^. In order to reach a high degree of saturation, 27 million AN3-12 mouse embryonic stem cells (mESCs) were used for mutagenesis. This was followed by TM selection using a WT lethal dose (0.5 µg/ml) for three weeks (Figure 1A). 29 resistant clones were randomly selected and picked to grow isogenic mutant lines. Whole exome sequencing was done with four clones, which showed strong TM resistance (Figure 1-figure supplement 1A). Two clones revealed independent missense mutations in the *Amdhd2* coding sequence (Figure 1-figure supplement 1B). A second round of whole exome sequencing of the remaining 25 clones revealed in total 11 independent amino acid substitutions at 10 distinct positions in *Amdhd2* (38% of sequenced clones) (Figure 1B, Figure 1-figure supplement 1B). Surprisingly we did not identify any mutations in the HBP’s rate limiting enzymes *Gfat1* or *Gfat2*. To confirm *Amdhd2* as the resistance-causing gene we generated *Amdhd2* K.O. mutants in WT AN3-12 cells using CRISPR/Cas9. We generated and validated a specific AMDHD2 antibody, which confirmed a successful K.O. of AMDHD2 (Figure 1C). Diploid *Amdhd2* K.O. cells showed significant TM resistance compared to WT cells, confirming AMDHD2 loss-of-function as causal for TM resistance (Figure 1D,E, Figure 1-figure supplement 1C). AMDHD2 is an amidohydrolase that plays a potential role in the HBP by catalyzing the deacetylation of GlcNAc-6P in the “reverse” direction of the pathway^20^. However, a role of AMDHD2 in modulating cellular UDP-GlcNAc levels has not been recognized before. Therefore, we hypothesized that AMDHD2 loss-of-function might increase UDP-GlcNAc levels leading to TM resistance (Figure 1F). To test this, we measured UDP-GlcNAc levels via ionic chromatography/mass spectrometry (IC-MS) and indeed AMDHD2 K.O. mutants showed significantly increased UDP-GlcNAc concentrations (Figure 1G, Figure 1-figure supplement 1D), indicating that the TM resistance is mediated by elevated HBP product availability due to reduced catabolism of GlcNAc-6P.

**Fig. 1:**
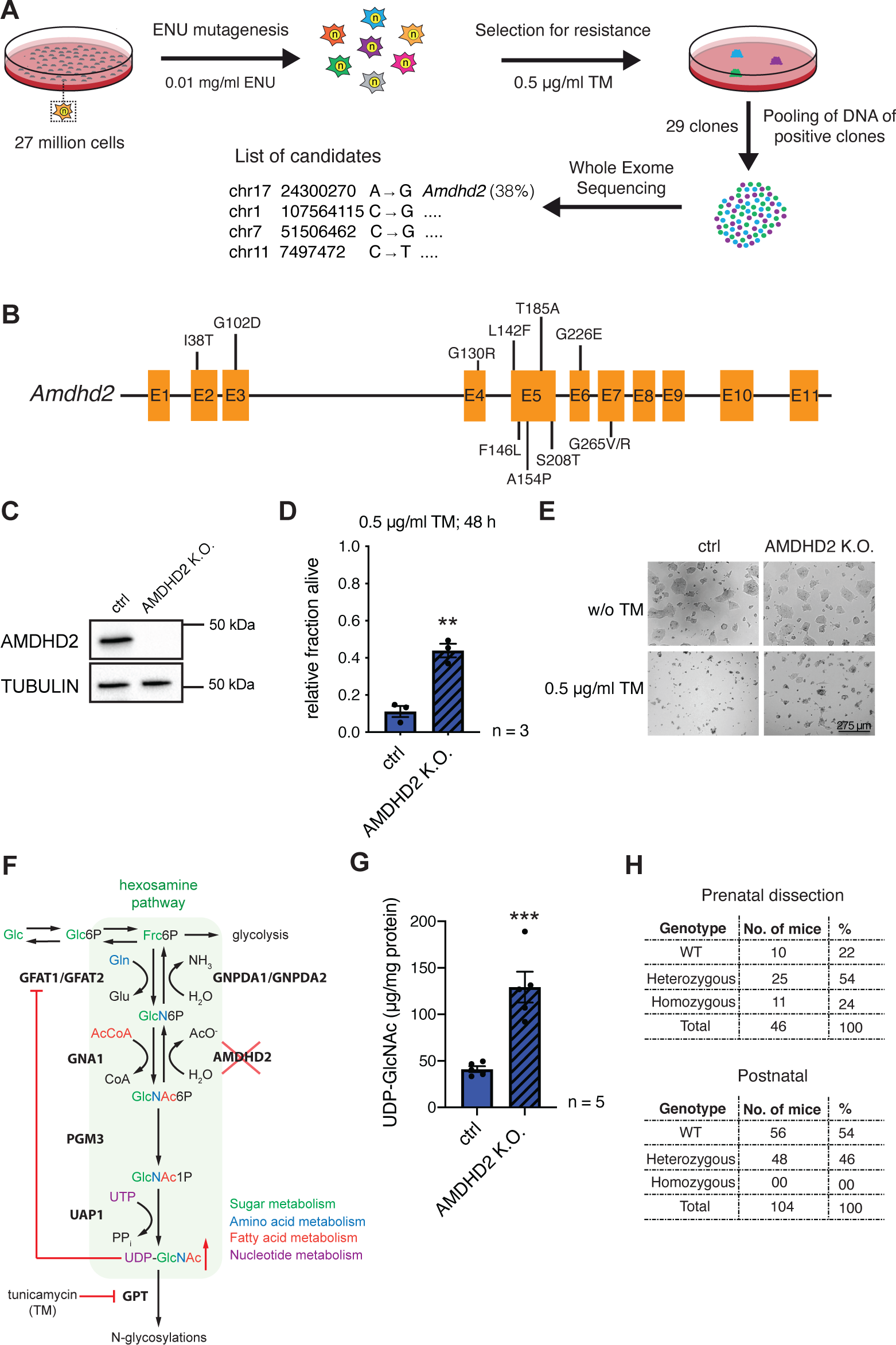
Chemical mutagenesis screen for tunicamycin resistance in mESCs identifies AMDHD2. **(A)** Schematic representation of experimental workflow for TM resistance screen using ENU mutagenesis in combination with whole exome sequencing. **(B)** Schematic representation of the mouse *Amdhd2* locus. Amino acid substitutions identified in the screen are highlighted. **(C)** Western blot analysis of CRISPR/Cas9 generated AMDHD2 K.O. AN3-12 mESCs compared to wildtype cells (ctrl). **(D)** Cell viability (XTT assay) of WT and AMDHD2 K.O. AN3-12 cells treated with 0.5 µg/ml TM for 48h (mean ± SEM, n=3, ** p<0.01, unpaired t-test). **(E)** Representative images of WT and AMDHD2 K.O. AN3-12 cells treated with 0.5 µg/ml TM for 48h or respective control. Scale bar, 275 µm**. (F)** Schematic overview of the hexosamine pathway (green box). The intermediate Frc6P from glycolysis is converted to UDP-GlcNAc, which is a precursor for glycosylation reactions. The enzymes are glutamine fructose-6-phosphate amidotransferase (GFAT1/2), glucosamine-6-phosphate N-acetyltransferase (GNA1), phosphoglucomutase (PGM3), UDP-N-acetylglucosamine pyrophosphorylase (UAP1), glucosamine-6-phosphate deaminase (GNPDA1/-), N-acetylglucosamine deacetylase (AMDHD2) and UDP- GlcNAc:dolichylphosphate GlcNAc-1-phosphotransferase (GPT). Red line indicates negative feedback inhibition of GFAT by UDP-GlcNAc. Formation of N- glycosylation is inhibited by tunicamycin (TM). **(G)** IC-MS analysis of UDP-GlcNAc levels of AMDHD2 K.O. compared to WT AN3-12 mESCs (mean ± SEM, n=5, *** p<0.001, unpaired t-test). **(H)** Genotyping results for the AMDHD2 deletion in dissected (E7-8) embryos and weaned mice. Figure supplements are available in Figure 1-figure supplements 1 and 2. Source data for this figure are available in Figure 1-source data 1.

To better understand the physiological consequences of HBP activation through AMDHD2 regulation, we disrupted the *Amdhd2* locus to generate a K.O. mouse (Figure 1-figure supplement 2A-C). Although the *Amdhd2* mutation distributed in Mendelian ratios in the offspring, no viable homozygous *Amdhd2* K.O. pups were weaned (Figure 1H), indicating a recessive mutation. Heterozygous animals however, did not show any obvious phenotype. Homozygous *Amdhd2* K.O. embryos showed early embryonic lethality, indicating an essential function of AMDHD2 during development. Taken together, we identified AMDHD2 as novel regulator of the HBP important in mESCs and for embryonic development.

### Structural and biochemical characterization of human AMDHD2

Until now, no structure of eukaryotic AMDHD2 was available and functional properties of human AMDHD2 remain largely unexplored. Therefore, we performed a structural and a biochemical characterization of human AMDHD2. Initial apo AMDHD2 crystals diffracted poorly and no structure could be solved. Based on homology to bacterial N-acetylglucosamine-6-phosphate deacetylase (NagA), human AMDHD2 is likely to bind a divalent cation in the active site, potentially stabilizing the protein and supporting co-crystallization. Consequently, we analyzed the stabilizing effect of several divalent cations. Addition of CoCl_2_, NiCl_2_, and ZnCl_2_ to the SEC buffer increased the thermal stability of AMDHD2 by 3-4°C (Figure 2A). Moreover, we tested the influence of CoCl_2_, NiCl_2_, and ZnCl_2_ on the deacetylase activity of AMDHD2. For that purpose, the metal co-factor of AMDHD2 was first removed by incubation with EDTA and then CoCl_2_, NiCl_2_, or ZnCl_2_ were added back. Addition of MgCl_2_ served as negative control, while an untreated AMDHD2 was used as positive control. Both CoCl_2_ and ZnCl_2_ restored and even increased the AMDHD2 activity (Figure 2B). Thus, Co^2+^ or Zn^2+^ might be the metal co-factor in human AMDHD2. We next tested co-crystallization of AMDHD2 with ZnCl_2_ or CoCl_2_. While no crystals formed in the presence of CoCl_2_, the co-crystallization with ZnCl_2_ yield needle clusters in several conditions. Optimized crystals diffracted to a resolution limit of 1.84 Å (AMDHD2 + Zn) or 1.90 Å (AMDHD2 + Zn + GlcN6P). The data collection and refinement statistics are summarized in Table 1. Human AMDHD2 is organized in two domains, a deacetylase domain responsible for the conversion of GlcNAc-6P into GlcN6P and a second small domain with unknown function (DUF) (Figure 2C, Figure 2- figure supplement 1). Residues from both the N-terminus and the C-terminus contribute to the DUF domain. The structure of AMDHD2 was almost completely modeled into the electron density map except for some N-terminal (1-5) and C-terminal residues (407-409). In the asymmetric unit, AMDHD2 forms a dimer through direct interactions of the deacetylase domains with an interface of 1117 Å^2^ and this dimeric assembly was judged as biological relevant by the EPPIC server^21^. Although the dimer is formed by a rather small interface, this conformation is supported by the crystallographic B-factors, which show low values at the interface, indicating a mutual stabilization (Figure 2-figure supplement 2). While AMDHD2 eluted as a pure monomer during SEC (Figure 2- figure supplement 3A), dynamic light scattering (DLS) measurements confirmed AMDHD2 dimers in solution (Figure 2-figure supplement 3B). These findings indicate that dimer formation might not be very stable. A comparison between both monomers from the dimer in the crystal revealed no major structural differences between monomer A and monomer B (Figure 2-figure supplement 4). The structure of the deacetylase domain showed a TIM (triosephosphate isomerase) barrel-like fold (Figure 2D). A typical TIM-barrel has eight alternating β-strands and α-helices forming a barrel shape where the parallel β-sheet builds the core that is surrounded by the α-helices. In AMDHD2, the eight alternating β-strands/α-helices are interrupted after eight β-strands and seven α-helices by an insertion of three antiparallel β-strands (β15-β17), which form an additional β-sheet close to the active site (Figure 2C, Figure 2-figure supplement 5). In monomer B, this β-sheet shows the highest crystallographic B-factors within the structure (Figure 2-figure supplement 2), indicating high flexibility and suggesting a functional role as a lid to the active site. The DUF-domain consists of two β-sheets, which are composed of three or six antiparallel β-strands each, and two small α-helices (Figure 2D). Together, these β-sheets form a β-sandwich. A superposition of the Zn-bound and the GlcN6P- and Zn-bound structures of AMDHD2 indicated no structural changes by the binding of the product (Figure 2- figure supplement 6). Residues from both monomers contribute to GlcN6P- binding (Figure 2E, Figure 2-figure supplement 7A). The phosphate group of the sugar is interacting via hydrogen bonds with Asn235 and Ala236, as well as ionic interactions to His242* and Arg243* of the other monomer (Figure 2E, Figure 2- figure supplement 7A,B). GlcN6P binding is further mediated by hydrogen bonds between the hydroxyl groups of GlcN6P with Ala154 and His272. The catalytic Zn ion is coordinated via electrostatic interactions with Glu143, His211, His232, and two water molecules, which in turn are stabilized by interactions with GlcN6P and several amino acid side chains including Asp294 that might, based on the homology to bacterial NagA, act as the catalytic base^22^ (Figure 2E, Figure 2- figure supplement 7A,B). We confirmed the presence of a single Zn ion in the human AMDHD2 active site by measuring an anomalous signal at the Zn-K edge (Figure 2F). Given the conservation of all functional residues (Figure 2-figure supplement 8), the human AMDHD2 reaction mechanism is likely to be very similar to the proposed mechanism for the enzyme from *E. coli* ^22^. In summary, our data show that human AMDHD2 is an obligate dimeric protein and carries a single catalytic Zn ion in the active center.

**Fig. 2:**
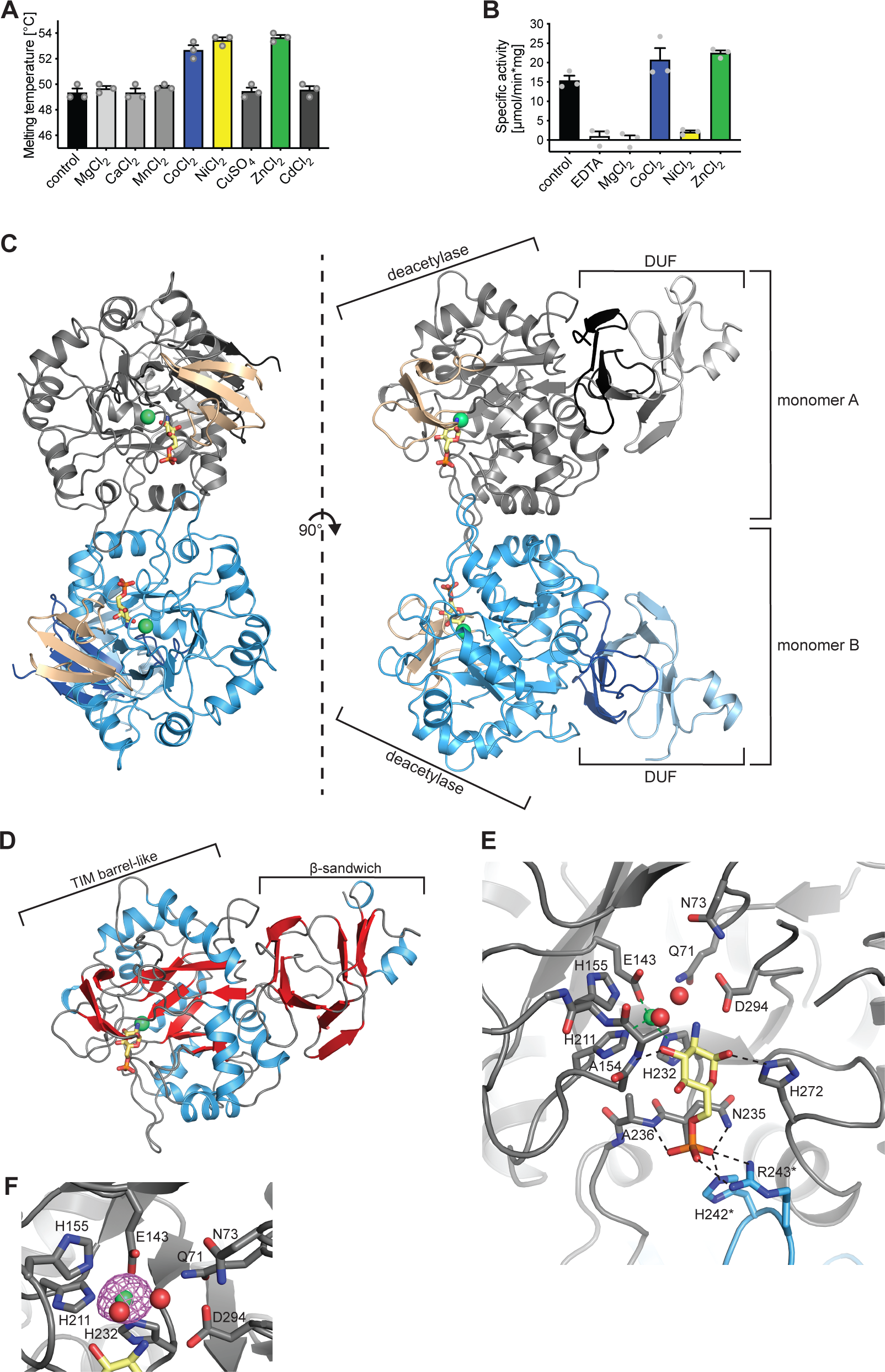
Structural and biochemical characterization of human AMDHD2. **(A)** Influence of divalent addition (10 µM) on the stability of AMDHD2 in SEC buffer in thermal shift assays (mean + SEM, n=3). **(B)** Deacetylase activity of AMDHD2 in the presence of EDTA and several indicated divalents (mean + SEM, n=3). **(C)** Overview of the human AMDHD2 dimer in cartoon representation. Monomer A is colored in gray and monomer B in blue. The two deacetylase domains are interacting with each other. The DUF domain is formed by residues of the N-terminus (light gray, light blue) and residues of the C-terminus (black, dark blue). GlcN6P (yellow sticks), Zn^2+^ (green sphere), and the putative active site lid (wheat) are highlighted. **(D)** Domains and secondary structure elements within one AMDHD2 monomer. The deacetylase domain (left) shows a TIM barrel-like fold, while the small DUF domain (right) is composed of a β-sandwich fold. α-helices are colored in blue, β-strands in red, and loops in gray. GlcN6P (yellow sticks) and Zn^2+^ (green sphere) are highlighted. **(E)** Close-up view of the active site in cartoon representation. Residues involved in ligand binding or catalysis are highlighted as sticks, as well as GlcN6P (yellow sticks), Zn^2+^ (green sphere) and two water molecules (red spheres). The GlcN6P binding site is formed by two deacetylase domains. Black dashed lines indicate key interactions to GlcN6P and green dashed lines the coordination of Zn^2+.^ **(F)** Anomalous map of Zn^2+^ with a contour level of 5.0 RMSD (violet). Figure supplements are available in Figure 2-figure supplements 1-8. Source data for this figure are available in Figure 2-source data 1.

**Table 1.** Data collection and refinement statistics of human AMDHD2.

|  | **AMDHD2**<br>**+ Zn + GlcN6P** | **AMDHD2**<br>**+ Zn** |
| --- | --- | --- |
| **Wavelength (Å)** | 1.00 | 1.00 |
| **Resolution range (Å)** | 45.71 - 1.90<br>(1.97 - 1.90) | 48.21 - 1.84<br>(1.90 - 1.84) |
| **Space group** | P 2<sub>1</sub> 2<sub>1</sub> 2<sub>1</sub> | P 2<sub>1</sub> 2<sub>1</sub> 2<sub>1</sub> |
| **a, b, c (Å)** | 63.3 161.4 86.6 | 61.8 84.3 154.2 |
| **α, β, γ (°)** | 90 90 90 | 90 90 90 |
| **Total reflections** | 428727 (42693) | 468961 (46539) |
| **Unique reflections** | 70760 (6907) | 71036 (6953) |
| **Multiplicity** | 6.1 (6.2) | 6.6 (6.7) |
| **Completeness (%)** | 99.7 (98.3) | 99.9 (99.2) |
| **Mean I/sigma(I)** | 11.46 (1.16) | 12.53 (1.06) |
| **Wilson B-factor** | 34.7 | 29.7 |
| **R<sub>merge</sub> (%)** | 9.5 (140.4) | 10.2 (150.6) |
| **R<sub>meas</sub> (%)** | 10.4 (153.3) | 11.0 (163.3) |
| **R<sub>pim</sub> (%)** | 4.2 (60.9) | 4.3 (62.5) |
| **CC<sub>1/2</sub> (%)** | 99.9 (49.4) | 99.9 (49.9) |
| **CC\* (%)** | 100 (81.3) | 100 (81.6) |
| **Reflections used in refinement** | 70751 (6906) | 71024 (6952) |
| **Reflections used for R-free** | 1980 (194) | 1992 (195) |
| **R<sub>work</sub> (%)** | 18.5 (31.7) | 18.2 (31.2) |
| **R<sub>free</sub> (%)** | 21.3 (29.6) | 20.6 (33.6) |
| **CC<sub>work</sub> (%)** | 96.6 (73.4) | 96.6 (75.7) |
| **CC<sub>free</sub> (%)** | 94.4 (73.1) | 95.4 (72.8) |
| **Number of non-hydrogen atoms** | 6361 | 6331 |
| **macromolecules** | 5997 | 5999 |
| **ligands** | 34 | 14 |
| **solvent** | 330 | 318 |
| **Protein residues** | 801 | 798 |
| **RMS (bonds) (Å)** | 0.005 | 0.004 |
| **RMS (angles) (°)** | 0.69 | 0.70 |
| **Ramachandran favored (%)** | 96.9 | 97.1 |
| **Ramachandran allowed (%)** | 2.9 | 2.7 |
| **Ramachandran outliers (%)** | 0.25 | 0.25 |
| **Rotamer outliers (%)** | 0.32 | 0.32 |
| **Clashscore** | 0.67 | 0.92 |
| **Average B-factor** | 43.59 | 37.67 |
| **macromolecules** | 43.81 | 37.76 |
| **ligands** | 40.19 | 41.98 |
| **solvent** | 40.10 | 35.72 |
| **Number of TLS groups** | 4 | 4 |
| **PDB ID** | 7NUT | 7NUU |

Statistics for the highest-resolution shell are shown in parentheses.

### Characterization of AMDHD2 loss-of-function mutants

We next characterized the eleven AMDHD2 substitutions from our screen and the putative active site mutant D294A to understand how they might affect the function of AMDHD2. Many AMDHD2 variants were soluble upon bacterial expression, including F146L, A154P, T185A, S208T, and D294A (Figure 3A). These substitutions are located close to the active site of AMDHD2 (Figure 3B) and Ala154 is involved in ligand binding by donating an H-bond via its main chain NH group to the 3-OH group of the sugar (Figure 2E, Figure 2-figure supplement 7A,B). In contrast, no soluble expression could be achieved for AMDHD2 G102D, G130R, G226E, and G265V (Figure 3A). The substitution of the small, flexible glycine by charged and/or bigger residues are likely to be incompatible with the proper tertiary structure and/or the folding process, thus resulting in insoluble AMDHD2 protein variants. The effect of the L142F mutation was even more severe as the substitution of Leu142 by the bigger phenylalanine resulted in AMDHD2 fragmentation (Figure 3A). Also, the I38T and G265R substitutions reduced soluble expression, indicating disturbed protein folding. We next tested the consequences of the I38T, T185A, G265R, and D294A substitutions on AMDHD2 activity. AMDHD2 T185A showed reduced activity and no activity was detected for G265R and D294A, while the third substitution, I38T, remained active (Figure 3C). This result indicates a functional role of Asp294 in the catalytic mechanism. It is very likely to act as catalytic base that is together with the metal ion activating the nucleophilic water molecule and later protonating the leaving group^22^. Moreover, the I38T substitution is the only identified mutation from the screen that is located in the DUF domain of AMDHD2. It reduced bacterial AMDHD2 expression yields, suggesting impaired protein folding. This is likely to result in a loss-of-function in vivo, while the purified and soluble protein is active. Taken together, the structural and biochemical characterization of AMDHD2 revealed that loss-of-function and subsequent HBP activation resulted from folding defects in AMDHD2 or it was caused by a loss of catalytic activity.

**Fig. 3:**
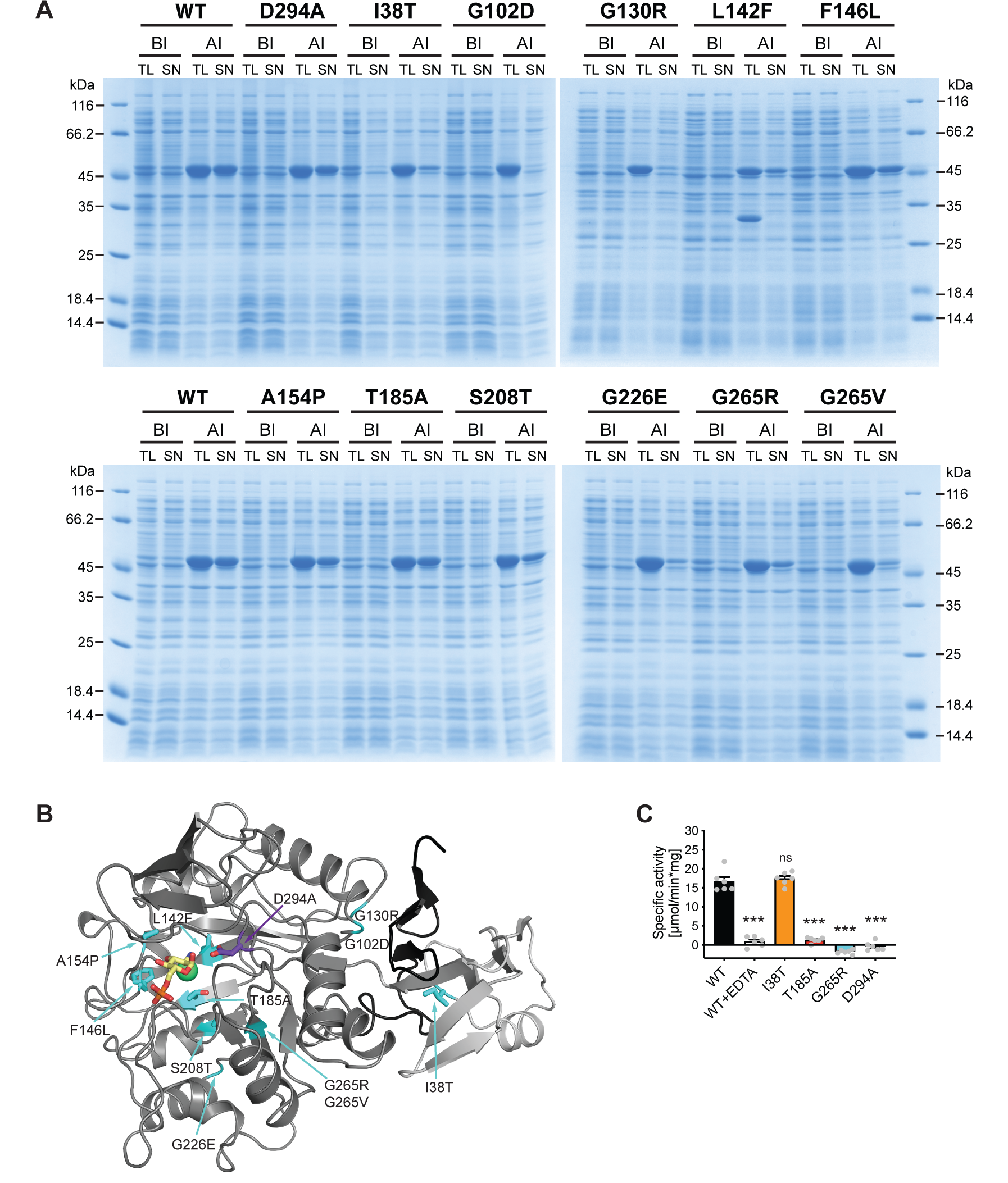
Characterization of AMDHD2 loss-of-function mutants. **(A)** SDS-gels stained with Coomassie brilliant blue of a representative bacterial test expression of the human AMDHD2 variants. The experiment was repeated three times with similar results. BI: before induction, AI: after induction, TL: total lysate, SN: soluble fraction/ supernatant. A band corresponding to the molecular weight of human AMDHD2 with His_6_-tag (46 kDa) was present in all total lysates after induction. **(B)** Overview of the position of the potential loss-of-function mutations in human AMDHD2 in cartoon representation. GlcN6P (yellow sticks), the metal co-factor (green spheres), the active site Asp294 (violet sticks), and the eleven putative loss-of-function mutations (cyan sticks) are highlighted. **(C)** Deacetylase activity of wild type (WT) and mutant human AMDHD2 (mean + SEM, n=6, *** p<0.0001 versus wild type, one-way ANOVA). Source data for this figure are available in Figure 3-source data 1.

### AMDHD2 limits HBP activity when GFAT2 replaces GFAT1 as the first enzyme

Having established that a loss of AMDHD2 function results in HBP activation, we were wondering about the role of the HBP’s rate limiting enzyme GFAT1. Under normal conditions, GFAT1 is constantly feedback inhibited by UDP-GlcNAc, crucially limiting HBP activity. A gain-of-function substitution in GFAT1 (G451E), however, increased HBP flux in nematodes and in murine cells, demonstrating a high degree of conservation^16^. In AN3-12 cells, the G451E gain-of-function substitution, introduced into the genomic locus by CRISPR/Cas9, as well as a *Gfat1* K.O. did not affect UDP-GlcNAc levels (Figure 4-figure supplement 1). While *Gfat1* is widely expressed across cell types, it is known that in some tissues *Gfat2* is the predominantly expressed paralog^7^. Since loss of GFAT1 did not affect HBP activity, we hypothesized that GFAT2 instead of GFAT1 might control metabolite entry into the HBP in AN3-12 mESCs. Indeed, *Gfat2* mRNA was abundantly expressed in AN3-12 cells, while expression levels of *Gfat1* were comparatively low (Figure 4A). Next, we performed WB analysis using defined amounts of pure purified human GFAT as standards to compare absolute GFAT abundance. GFAT2 was found abundantly expressed, while GFAT1 was difficult to detect in AN3-12 mESCs (Figure 4B). E14 mESCs likewise showed predominant GFAT2 expression and low GFAT1 abundance. In contrast, mouse neuronal N2a cells as well as muscle precursor C2C12 myoblasts showed predominant GFAT1 expression and GFAT2 was virtually undetectable. These data suggest a HBP configuration characterized by a high GFAT2/GFAT1 ratio in mESCs.

**Fig. 4:**
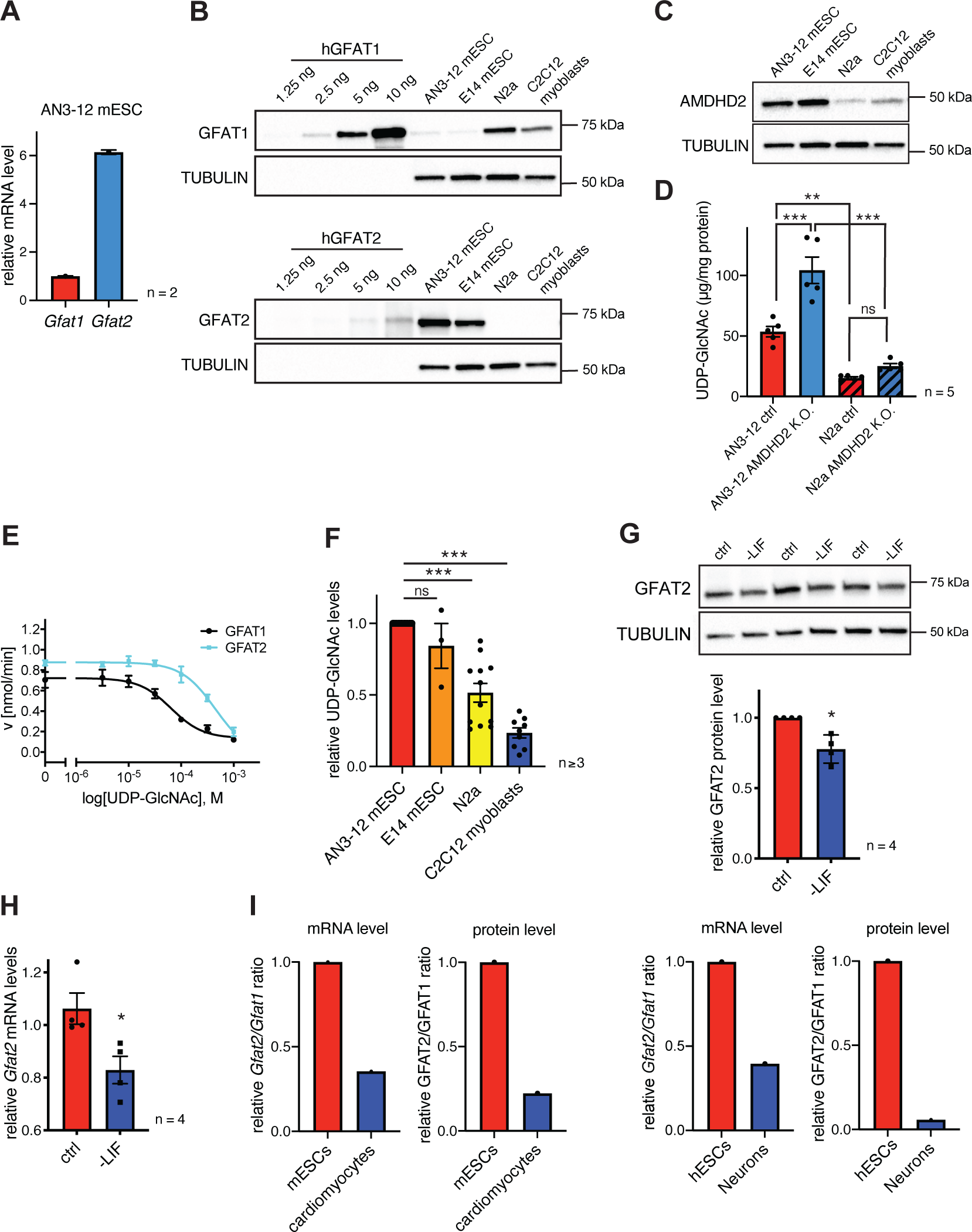
AMDHD2 limits HBP activity when GFAT2 replaces GFAT1 as the first enzyme. **(A)** Relative *Gfat1* and *Gfat2* mRNA levels (qPCR) in WT AN3-12 cells (mean + SEM, n=2). **(B)** Western blot analysis of indicated amounts of purified human GFAT1 and GFAT2 protein as standards to compare GFAT abundance in cell lysates of indicated cell lines. **(C)** Western blot analysis of AMDHD2 in indicated cell lines. **(D)** IC-MS analysis of UDP-GlcNAc levels in WT and AMDHD2 K.O. AN3-12 mESCs and N2a cells (mean ± SEM, n=5, *** p<0.001, one-way ANOVA). **(E)** Representative UDP-GlcNAc dose-response assay with hGFAT1 (black circle) and hGFAT2 (teal square) (mean ± SD, n=3). **(F)** Relative UDP-GlcNAc levels in indicated cell lines measured by IC-MS. Levels are normalized to those in AN3-12 mESCs (mean ± SEM, n≥3, *** p<0.001, one-way ANOVA) **(G)** Western blot analysis and quantification (mean ± SD, n=4, * p<0.05, unpaired t-test) of GFAT2 in WT AN3-12 control cells and upon partial differentiation by a 5-day LIF removal. **(H)** Relative *Gfat2* mRNA level (qPCR) in WT AN3-12 cells and upon partial differentiation by a 5-day LIF removal (mean ± SEM, n=4, * p<0.05, unpaired t-test). **(I)** Relative *Gfat2/Gfat1* mRNA and GFAT2/GFAT1 protein ratios in mouse or human ESCs and their differentiated counterparts as indicated. Figure supplements are available in Figure 4-figure supplements 1-4. Source data for this figure are available in Figure 4-source data 1.

To further investigate the possibility of ESC specific HBP regulation, we next checked AMDHD2 levels in mESCs and compared them to cells using GFAT1 as the predominant first HBP enzyme. Consistent with GFAT2 levels, AMDHD2 protein abundance was higher in mESCs compared to N2a and C2C12 cells (Figure 4C). Moreover, the K.O. of AMDHD2 in AN3-12 mESCs resulted in a drastic elevation of UDP-GlcNAc levels, while the loss of AMDHD2 in N2a cells had no significant effect (Figure 4D). This indicates that AMDHD2 was constitutively active in AN3-12 cells, while the catalysis of the reverse flux of the HBP by AMDHD2 seemed to be negligible in N2a cells. We therefore hypothesized that AMDHD2 plays a key role in the HBP when GFAT2 is its first enzyme instead of the more common GFAT1. Our previous data indicate that GFAT1 is under constant UDP-GlcNAc inhibition, sufficient for full suppression of GFAT1 activity^15^. We reasoned that higher UDP-GlcNAc levels in mESCs can only be achieved by differences in UDP-GlcNAc feedback inhibition between GFAT1 and GFAT2. To address this point, we generated recombinant human GFAT1 and GFAT2 with an internal His_6_tag and characterized the proteins in activity assays (Figure 4E, Table 2, Figure 4-figure supplement 2A,B). Kinetic measurements confirmed that both proteins were fully functional and revealed different substrate affinities of GFAT2 compared to GFAT1 (Table 2, Figure 4- figure supplement 2A,B). In a UDP-GlcNAc dose response assay, we found a significantly higher IC_50_ value for GFAT2 (367.3 -43.6/+49,5 µM) compared to GFAT1 (57.0 -8.3/+9.7 µM) (Figure 4E, Table 2). We conclude, first, that UDP-GlcNAc inhibition is weaker in GFAT2 compared to GFAT1 and, second, that AMDHD2 plays a crucial role in balancing GFAT2 mediated HBP flux. Consistent with lower feedback inhibition of GFAT2, UDP-GlcNAc levels and protein O-GlcNAc modification in AN3-12 mESCs were significantly higher than in N2a and C2C12 cells with a GFAT1-regulated HBP (Figure 4F, Figure 4-figure supplement 3).

**Table 2.** Kinetic parameters of human GFAT1 and GFAT2.

|  | L-Glu production |  |  | D-GlcN6P production |  |  | UDP-GlcNAc inhibition |
| --- | --- | --- | --- | --- | --- | --- | --- |
| | $K_m$ L-Gln<br>[mM] | $k_{cat}$<br>[s<sup>-1</sup>] | $k_{cat}/K_m$<br>[mM<sup>-1</sup> s<sup>-1</sup>] | $K_m$ Frc6P<br>[mM] | $k_{cat}$<br>[s<sup>-1</sup>] | $k_{cat}/K_m$<br>[mM<sup>-1</sup> s<sup>-1</sup>] | IC<sub>50</sub><br>[μM] |
| **GFAT-1** | 1.1 ± 0.19 | 3.6 ± 0.18 | 3.3 | 0.08 ± 0.01 | 1.7 ± 0.09 | 21.3 | 57.0<br>-8.3/+9.7 |
| **GFAT-2** | 0.5 ± 0.06 | 3.7 ± 0.10 | 7.4 | 0.29 ± 0.05 | 1.8 ± 0.09 | 6.2 | 367.3<br>-43.6/+49.5 |
| **Unpaired**<br>**t-test**<br>**(two-**<br>**sided)** | ** p=0.005 |  |  | ** p=0.0027 |  |  | *** p=0.0002 |

In a next step, we asked if differentiation of mESC might affect the HBP’s enzymatic configuration. For this, we removed leukemia inhibitory factor (LIF) from the medium, initiating differentiation^23^. LIF removal for five days resulted in partial differentiation of AN3-12 cells as indicated by a decrease of stem cell markers (Figure 4-figure supplement 4A). Of note, GFAT2 protein as well as *Gfat2* mRNA levels decreased significantly with LIF removal (Figure 4G,H). GFAT1 and AMDHD2 mRNA and protein levels did not change in this partial differentiation paradigm (Figure 4-figure supplement 4B,C). A decrease in the GFAT2/GFAT1 ratio upon differentiation was also observed in published datasets: relative GFAT2 mRNA and protein levels decrease during neuronal differentiation of human ESCs^24^ and during mESC differentiation in the cardiac lineage^25^ (Figure 4I).

Taken together, our data indicate a configuration of the HBP that is specific to mESCs, in which the rate limiting reaction is regulated by GFAT2 instead of GFAT1. Given the weak feedback inhibition of GFAT2 there is a need for balancing HBP metabolic flux and our data demonstrate that AMDHD2 fulfills this role in mESCs.

## Discussion

HBP activation increases cellular UDP-GlcNAc levels that allosterically protect from TM toxicity^3^. We used this knowledge to interrogate the HBP for additional regulators in a forward genetic TM resistance screen using haploid cells. Random chemical DNA mutagenesis at high saturation in haploid cells is a unique strategy to identify recessive mutations including those leading to single amino acid substitutions. Using this approach, we identified the N-acetylglucosamine-6- phosphate deacetylase AMDHD2 as a novel regulator of the mammalian HBP. We next solved the first crystal structure of human AMDHD2 and noted that resistance-associated substitutions disturb protein folding or cluster in the catalytic pocket, likely interfering with substrate binding or catalysis. Finally, we found that mESCs utilize GFAT2 for metabolite entry into the HBP instead of the more widely expressed GFAT1. GFAT2 is under considerably reduced UDP-GlcNAc feedback inhibition explaining why loss of AMDHD2 activity was sufficient for HBP activation without GFAT mutations.

Chemical mutagenesis-based screening in haploid cells represents a powerful and unique technique. This state-of-the-art approach allows to dissect the entire spectrum of mutations including loss-of-function, gain-of-function, and neomorph alleles^18^. The additional usage of haploid cells not only enables detection of dominant but also recessive mutations due to the lack of a remaining and interfering WT allele. Of note, identification of AMDHD2 as a novel regulator of the HBP was only possible in this specific setup since *Amdhd2* mutations are recessive as shown in the AMDHD2 K.O. mouse.

Besides the function as GlcNAc-6P deacetylase, AMDHD2 was shown to be involved in the degradation of N-glycolylneuraminic acid (Neu5Gc) in mice and in human cell culture^26, 27^. Nevertheless, mammalian AMDHD2 is rather unstudied and most knowledge is based on the bacterial homolog NagA. NagA catalyzes the deacetylation reaction in the HBP, contributing to recycling of cell wall components such as GlcNAc. Since breakdown of GlcNAc can be used as an energy source by bacteria and fungi, NagA plays a crucial role in their energy metabolism^28–31^. For this reason, HBP enzymes are attractive selective targets for antifungal and antibiotic drugs^32–35^. While catabolism of aminosugars connects GlcNAc with glycolysis, AMDHD2 had not been implicated in a regulatory role of the HBP and cellular UDP-GlcNAc homeostasis.

After identification of AMDHD2 as a key modulator of the mammalian HBP, we structurally and biochemically characterized human AMDHD2. We solved the structure of human AMDHD2, the first reported eukaryotic structure of an AMDHD2 homolog. AMDHD2 is a dimeric enzyme and residues from both monomers contribute to ligand binding in the active site, while the residues important for catalysis originate from one monomer. Thus, monomeric AMDHD2 might be active, with weaker substrate affinity or specificity. The oligomeric state of AMDHD2 is therefore a plausible target to modulate its catalytic properties. Furthermore, the involvement of both monomers in ligand binding suggests ligand-induced dimerization *in vivo*. Bacterial NagAs are reported to use N-acetylgalactosamine-6-phosphate (GalNAc-6P), N-acetylmannosamine-6- phosphate (ManNAc-6P), and N-acetylglucosamine-6-sulphate (GlcNAc-6S) as substrates, albeit with increased K_m_ values^22, 36^. The high structural conservation of the side chains interacting with the sugar’s C4 for GalNAc-6P, C2 for ManNAc-6P or the phosphate group indicate that human AMDHD2 might catalyze the deacetylation of several N-acetyl amino sugars as well.

We showed that the mutations identified in the screen cause a loss-of-function in human AMDHD2 by disrupting its folding or activity (Figure 3). AMDHD2 is composed of a deacetylase domain and a small domain with unknown function (DUF). We identified only one mutation, I38T, within the DUF domain and this mutant showed diminished expression yields and low solubility, potentially explaining the loss-of-function. Nonetheless, the soluble fraction of AMDHD2 I38T was as active as wild type AMDHD2 in activity assays, indicating that the DUF domain might be dispensable for catalysis.

Further characterizing the HBP, we noticed a surprising configuration of HBP enzymes in AN3-12 and E14 mESCs. While N2a cells and C2C12 myoblasts rely on GFAT1 as the key HBP enzyme, the mESCs use GFAT2 that is abundantly expressed. Consistently, genetic manipulation of GFAT1 did not show any effect on UDP-GlcNAc levels in AN3-12 mESCs, while introducing the G451E gain of function mutation in GFAT1 of N2a cells leads to the previously reported boost of HBP activity^16^. Additionally, AMDHD2 abundance was higher in mESCs (Figure 4C). In accordance, the AMDHD2 K.O. in AN3-12 mESCs massively elevated UDP-GlcNAc levels, while the loss of AMDHD2 in N2a cells had no significant impact. Under physiological conditions, GFAT1 is strongly inhibited by UDP-GlcNAc^15^. In this scenario, as is the case in N2a cells, loss of the reverse flux by AMDHD2 K.O. showed no drastic effect on UDP-GlcNAc levels. Moreover, we showed that GFAT2 has altered substrate affinities and is less susceptible to UDP-GlcNAc feedback inhibition. N- or C-terminal tags in GFAT disturb the catalytic function, therefore the GFAT preparations used here carry an internal tag for purification at a position that is reported not to interfere with the kinetic properties of GFAT1^37^. Studies with other tagging strategies reported only a weak inhibition of GFAT2 by UDP-GlcNAc^38, 39^. In contrast, we demonstrate that GFAT2 can be fully inhibited by UDP-GlcNAc, to an extent similar to GFAT1. However, this required approximately 6-fold higher UDP-GlcNAc concentrations. Overall, our data suggest that GFAT1 is sufficiently regulated by feedback inhibition to determine HBP flux. Cells using GFAT2 in the HBP, in contrast, rely on AMDHD2 to balance forward and reverse flux in the HBP.

This HBP configuration might be a general adaptation of mESCs as we could show similar results in E14 mESCs. Differentiation might then switch GFAT expression and indeed partial differentiation of AN3-12 cells by LIF removal induced a significant decrease in GFAT2 levels. GFAT1 and AMDHD2 levels were not affected likely due to the early differentiation state. Analysis of published data confirmed that GFAT2 is highly expressed in human ESCs and mESCs, and abundance decreased during neuronal or myocyte differentiation. Consistent with these findings, intestinal stem cells in *Drosophila melanogaster* likewise express GFAT2^40^. One potential consequence of this metabolic adaptation in mESCs is a higher baseline UDP-GlcNAc concentration compared to cells that use GFAT1 to control the HBP. This increase in UDP-GlcNAc concentration might affect downstream PTMs, which in turn can influence cell signaling. In particular, O-GlcNAc modifications already have been linked to stemness and pluripotency^41, 42^. Indeed, we detected increased O-GlcNAc levels in mESCs compared to cells utilizing GFAT1 in the HBP. Additional significance of an ESC-specific HBP configuration might come from an adaptation to their special nutrient and energy requirements. ESCs show a specialized metabolic profile that likely affect the concentrations of GFAT substrates^43^. The kinetic properties of GFAT2 might reflect an adaption to substrate availability in ESCs.

Taken together, we identify AMDHD2 as an essential gene and describe a cell type specific role of AMDHD2 acting in tandem with GFAT2 to regulate the HBP. Tuning HBP metabolic activity is relevant in cellular stress resistance, oncogenic transformation and growth, and in longevity. Our work advances the understanding of HBP control and provides specific means to beneficially affect these processes in the future.

## Methods

### Cell lines and culture conditions

AN3-12 mouse embryonic haploid stem cells were cultured as previously described^17^. In brief, DMEM high glucose (Sigma-Aldrich) was supplemented with glutamine, fetal bovine serum (15%), penicillin/streptomycin, non-essential amino acids, sodium pyruvate (all Thermo Fisher Scientific, Waltham, Massachusetts), β-mercaptoethanol and LIF (both Merck Millipore) and used to culture cells at 37°C in 5% CO_2_ on non-coated tissue culture plates. For partial differentiation of AN3-12 cells, cells were seeded at a density of 2000-3000 cells/6-well and incubated for 5 days in medium without LIF.

N2a mouse neuroblastoma cells (ATCC) and C2C12 (ATCC) cells were cultured in DMEM containing 4.5 g/l glucose (Gibco) supplemented with 10% fetal bovine serum (Gibco) and penicillin/streptomycin at 37°C in 5% CO_2_.

### Cell sorting

To maintain a haploid cell population cells were stained with 10 μg/ml Hoechst 33342 (Thermo Fisher Scientific) for 30 min at 37°C. To exclude dead cells propidium iodide (Sigma-Aldrich) staining was added. Cells were sorted for DNA content on a FACSAria Fusion sorter and flow profiles were recorded with the FACSDiva software (BD Franklin Lakes).

### Cell viability assay (XTT)

Relative cell viability was assessed using the XTT cell proliferation Kit II (Roche Diagnostics) according to the manufacturer’s instructions. Tunicamycin treatments were performed for 48 hours, starting 24 hours after cell seeding. XTT turnover was normalized to corresponding untreated control cells.

### Mutagenesis screen, exome sequencing, and analysis

The screening procedure and the data analysis were extensively described previously^18^. In brief, AN3-12 mouse embryonic haploid stem cells were mutagenized with 0.01 mg/ml Ethylnitrosourea for 2h at room temperature prior to drug selection starting 24 hours post mutagenesis using 0.5 μg/ml tunicamycin (Sigma-Aldrich). After 21 days of drug selection, resistant clones were isolated and subjected to tunicamycin cytotoxicity assays and gDNA extraction using the Gentra Puregene Tissue Kit (Qiagen). Paired end, 150 bp whole exome sequencing was performed on an Illumina Novaseq 6000 instrument after precapture-barcoding and exome capture with the Agilent SureSelect Mouse All Exon kit. For data analysis, raw reads were aligned to the reference genome mm9. Variants were identified and annotated using GATK (v.3.4.46) and snpEff (v.4.2). Tunicamycin resistance causing alterations were identified by allelism only considering variants with moderate or high effect on protein and a read coverage > 20.

### Gene editing and genotyping by Sanger sequencing

The specific GFAT1 G451E substitution as well as the K.O. of GFAT1 and AMDHD2 was engineered in AN3-12 cells (for the AMDHD2 K.O. also in N2a cells) using the CRISPR/Cas9 technology as described previously^44^. DNA template sequences for small guide RNAs were designed online (http://crispor.org, Supplementary Table 1), purchased from Sigma, and cloned into the Cas9-GFP expressing plasmid PX458 (Addgene #48138). Corresponding guide and Cas9 expressing plasmids were co-transfected with a single stranded DNA repair template (Integrated DNA technologies), using Lipofectamine 3000 (Thermo Fisher Scientific) according to manufacturer’s instructions. GFP positive cells were singled using FACSAria Fusion sorter and subjected to genotyping. DNA was extracted (DNA extraction solution, Epicentre Biotechnologies) and edited regions were specifically amplified by PCR (primers are listed in Supplementary Table 1). Sanger sequencing was performed at Eurofins Genomics GmbH (Ebersberg, Germany).

### RNA isolation and qPCR

Cells were collected in QIAzol (Qiagen) and snap frozen in liquid nitrogen. Samples were subjected to three freeze/thaw cycles (liquid nitrogen/ 37°Cwater bath) before addition of another half of the total QIAzol volume. After incubation for 5 min at RT, 200 μl chloroform were added per 1 ml QIAzol. Samples were vortexed, incubated for 2 min at RT, and centrifuged at 10.000 rpm and 4°C for 15 min. The aqueous phase was mixed with an equal volume of 70% ethanol and transferred to a RNeasy Mini spin column (Qiagen). The total RNA was isolated using the RNeasy Mini Kit (Qiagen) and cDNA was subsequently generated by iScript cDNA Synthesis Kit (BioRad). qPCR was performed with Power SYBR Green master mix (Applied Biosystems) on a ViiA 7 Real-Time PCR System (Applied Biosystems). GAPDH expression functioned as internal control. All used primers for qPCR analysis are listed in Supplementary Table 1.

### Anion exchange chromatography mass spectrometry (IC-MS) analysis of UDP-GlcNAc and UDP-GalNAc

Cells were subjected to methanol:acetonitrile:mili-Q ultrapure water (40:40:20 [v:v:v]) extraction. UDP-GlcNAc and UDP-GalNAc (UDP-HexNAc) concentrations were measured using IC-MS analysis. Extracted metabolites were re-suspended in 500 µl of Optima LC/MS grade water (Thermo Fisher Scientific) of which 100 µl were transferred to polypropylene autosampler vials (Chromatography Accessories Trott, Germany). The samples were analyzed using a Dionex ionchromatography system (ICS5000, Thermo Fisher Scientific) connected to a triple quadrupole MS (Waters, TQ). In brief, 10 µl of the metabolite extract were injected in full loop mode using an overfill factor of 3, onto a Dionex IonPac AS11-HC column (2 mm × 250 mm, 4 μm particle size, Thermo Scientific) equipped with a Dionex IonPac AG11-HC guard column (2 mm × 50 mm, 4 μm, Thermo Scientific). The column temperature was held at 30°C, while the auto sampler was set to 6°C. The metabolite separation was carried using a KOH gradient at a flow rate of 380 µl/min, applying the following gradient conditions: 0-8 min, 30-35 mM KOH; 8-12 min, 35−100 mM KOH; 12-15 min, 100 mM KOH, 15-15.1 min, 10 mM KOH. The column was re-equilibrated at 10 mM for 4 min. UDP-HexNAcs were detected using multiple reaction monitoring (MRM) mode with the following settings: capillary voltage 2.7 kV, desolvation temperature 550°C, desolvation gas flow 800 l/h, collision cell gas flow 0.15 ml/min. The transitions for UDP-GalNAc, as well as for UDP-GlcNAc were m/z 606 [M-H+]+ for the precursor mass and m/z 385 [M-H+]+ for the first and m/z 282 [M-H+]+ for the second transition mass. The cone voltage was set to 46V and the collision energy was set to 22V. UDP-GalNAc eluted at 10.48 min and UDP-GlcNAc eluted at 11.05 min. MS data analysis was performed using the TargetLynx Software (Version 4.1, Waters). Absolute compound concentrations were calculated from response curves of differently diluted authentic standards treated and extracted as the samples.

### Immunoblot analysis

Protein concentration of cell lysates was determined using the Pierce^TM^ BCA protein assay kit according to manufacturer’s instructions (ThermoFisher Scientific). Samples were adjusted in 5xLDS sample buffer containing 50 mM DTT. After boiling and a sonication step, equal protein amounts were subjected to SDS-PAGE and blotted on a nitrocellulose membrane using the Trans-Blot Turbo Transfer system (BioRad). All antibodies were used in 5% low-fat milk or 5% BSA in TBS-Tween. After incubation with HRP-conjugated secondary antibody, the blot was developed using ECL solution (Merck Millipore) on a ChemiDoc MP Imaging System (BioRad).

The following antibodies were used in this study: GFAT1 (rb, EPR4854, Abcam ab125069, 1:1000), GFAT2 (rb, Abcam, ab190966, 1:5000), O-Linked N-Acetylglucosamine Antibody (ms, clone RL2, MABS157, Merck, 1:1000), AMDHD2 (ms, S6 clone, in-house produced, 1:500), *α*-TUBULIN (ms, clone B-5- 1-2, Sigma T9026, 1:5000), rabbit IgG (gt, HRP-conjugated, G21234, Thermo Fisher,1:5000), and mouse IgG (gt, HRP-conjugated, G21040, Thermo Fisher, 1:5000).

### Generation of anti-AMDHD2 antibody

To generate monoclonal antibodies directed against AMDHD2, His-tagged human AMDHD2 was expressed in *Escherichia coli*, affinity purified, and used for immunization of eight-week-old male Balb/cJRj mice. The first immunization with 80 µg of recombinant protein was enhanced by Freund’s complete adjuvant; subsequent injections used 40 µg protein with Freund’s incomplete adjuvant. After multiple immunizations, the serum of the mice was tested for immunoreaction by enzyme-linked immunosorbent assay (ELISA) with the recombinant His-hAMDHD2 protein. In addition, the serum was used to stain immunoblots with lysates of HEK293T cells overexpressing FLAG-HA- hAMDHD2. After this positive testing, cells from the popliteal lymph node were fused with mouse myeloma SP2/0 cells by a standard fusion protocol. Monoclonal hybridoma lines were characterized, expanded, and subcloned according to standard procedures^45^. Initial screening of clones was performed by ELISA with recombinant His-AMDHD2 protein and immunoblots using FLAG-HA- hAMDHD2 overexpressed in HEK293T cells. Isotyping of selected clones was performed with Pierce Rapid Isotyping Kit (Thermo Scientific, #26179). Final validation of antibody specificity was done by immunoblots of WT N2a cells compared to cells overexpressing FLAG-HA-hAMDHD2 and AMDHD2 K.O. cells.

### Expression and purification of human AMDHD2

A pET28a(+)-AMDHD2 plasmid was purchased from BioCat (Heidelberg, Germany), where human AMDHD2 isoform 1 was integrated in pET28a(+) using NdeI and HindIII restriction sites. This vector was used to recombinantly express human AMDHD2 isoform 1 with N-terminal His_6_tag and a thrombin cleavage site under the control of the T7 promoter in BL21 (DE3) *E. coli.* LB cultures were incubated at 37°C and 180 rpm until an OD_600_ of 0.4-0.6 was reached. Then, protein expression was induced by addition of 0.5 mM isopropyl-β-D-1- thiogalactopyranosid (IPTG) and incubated for 20-22 h at 20°C and 180 rpm. Cultures were harvested and pellets stored at -80°C. The purification buffers were modified from Bergfeld *et al.*^26^. *E. coli* were lysed in 50 mM Tris/HCl pH 7.5, 100 mM NaCl, 20 mM imidazole, 1 mM Tris(2-carboxyethyl)phosphin (TCEP) with complete EDTA-free protease inhibitor cocktail (Roche) and 10 µg/ml DNAseI (Sigma) by sonication. The lysate was clarified by centrifugation and the supernatant loaded on Ni-NTA Superflow affinity resin (Qiagen). The resin was washed with wash buffer (50 mM Tris-HCl, 100 mM NaCl, 50 mM imidazole, 1 mM TCEP; pH 7.5) and the protein was eluted with wash buffer containing 250 mM imidazole. The His_6_-tag was proteolytically removed using 5 Units of thrombin (Sigma-Aldrich) per mg protein overnight at 4°C. AMDHD2 was further purified according to its size on a HiLoad™ 16/60 Superdex™ 200 prep grade prepacked column (GE Healthcare) using an ÄKTAprime chromatography system at 4°C with a SEC buffer containing 50 mM Tris-HCl, 100 mM NaCl, 1 mM TCEP, 5 % glycerol; pH 7.5.

### Site-directed mutagenesis

The AMDHD2 mutations were introduced into the pET28a(+)-AMDHD2 plasmid by site-directed mutagenesis as described previously^46^ (Mutagenesis primers are listed in Supplementary Table 1). This protocol was also used to integrate an internal His_6_-tag between Ser300 and Asp301 in human GFAT2 in the plasmid FLAG-HA-hGFAT2-pcDNA3.1 (pcDNA™3.1(+), ThermoFisher Scientific #V79020). This position is equivalent to the internal His_6_-tag in human GFAT1, which does not interfere with GFAT kinetic properties^37^. The GFAT2 gene with internal His_6_-tag was subsequently subcloned into the pFL vector for the generation of baculoviruses using XbaI and HindIII entry sites.

### Thermal shift assay

The thermal stability of AMDHD2 was analyzed by thermal shift (thermofluor) assays. For this purpose, the proteins were incubated with SYPRO orange dye (Sigma-Aldrich), which binds specifically to hydrophobic amino acids leading to an increased fluorescence at 610 nm when excited with a wavelength of 490 nm. The melting temperature is defined as the midpoint of temperature of the protein- unfolding transition^47^. This turning point of the melting curve was extracted from the derivative values of the RFU curve, where a turning point to the right is a minimum. The influence of several divalent cations on the thermal stability of AMDHD2 was tested. For this, the SEC buffer was supplemented with MgCl_2_, CaCl_2_, MnCl_2_, CoCl_2_, NiCl_2_, CuSO4, ZnCl_2_, or CdCl_2_ at a final concentration of 10 µM. The reaction mixtures were pipetted in white RT-PCR plates and contained 5 µl SYPRO orange dye (1:500 dilution in ddH_2_O) and 5-10 µg protein in a total volume of 50 µl. The plates were closed with optically clear tape and placed in a BioRad CFX-96 Real-Time PCR machine. The melting curves were measured at 1°C/min at the FRET channel in triplicate measurements and the data analyzed with CFX Manager™ (BioRad).

### GlcN6P production of AMDHD2

The deacetylase activity of AMDHD2 was determined by following the cleavage of the amide/peptide bond of GlcNAc6P at 205 nm in UV transparent 96 well microplates (F-bottom, Brand #781614). The assay mix contained 1 mM GlcNAc6P in 50 mM Tris-HCl pH 7.5 and was pre-warmed for 10 min at 37 °C in the plate reader. The reaction was started by adding 20 pmol AMDHD2 and was monitored several minutes at 37°C. The initial reaction rates (0-1 min) were determined by Excel (Microsoft) and the amount of consumed GlcNAc6P was calculated from a GlcNAc6P standard curve. All measurements were performed in duplicates. For the analysis of the impact of several divalent metal ions on the activity of AMDHD2, the protein was incubated for 10 min with 0.1 µM EDTA and afterwards 10 µM divalent was added to potentially restore activity.

### Human AMDHD2 crystallization and crystal soaking

Human AMDHD2 was co-crystallized with a 1.25x ratio (molar) of ZnCl_2_ at a concentration of 9 mg/ml in sitting-drops by vapor diffusion at 20°C. Intergrown crystal plates formed in the PACT *premier*™ HT-96 (Molecular Dimensions) screen in condition H5 with a reservoir solution containing 0.1 M bis-tris propane pH 8.5, 0.2 M sodium nitrate, and 20% (w/v) PEG3350. In an optimization screen, the concentration of PEG3350 was constant at 20 % (w/v), while the pH value of bis-tris propane and the concentration of sodium nitrate were varied. The drops were set up in 1.5 μl protein solution to 1.5 μl precipitant solution and 2 μl protein solution to 1 μl precipitant solution. Best crystals were obtained with a drop ratio of 2 μl protein solution to 1 μl precipitant solution at 0.1 M bis tris propane pH 8.25, 0.25 M sodium nitrate, and 20% (w/v) PEG3350. 5 mM GlcN6P in reservoir solution was soaked into the crystals for 2 to 24h. For crystal harvesting, the intergrown plates were separated with a needle and 15% glycerol was used as cryoprotectant.

### Data collection and refinement

X-ray diffraction measurements were performed at beamline P13 at PETRA III, DESY, Hamburg (Germany) and beamline X06SA at the Swiss Light Source, Paul Scherrer Institute, Villigen (Switzerland). The diffraction images were processed by XDS^48^. The structure of human AMDHD2 was determined by molecular replacement^49, 50^ with phenix.phaser^51, 52^ using the models of *B. subtilis* AMDHD2 (PDB 2VHL) as search model. The structures were further manually built using COOT^53^ and iterative refinement rounds were performed using phenix.refine^52^. The structure of GlcN6P soaked crystals was solved by molecular replacement using our human AMDHD2 structure as search model. Geometry restraints for GlcN6P was generated with phenix.elbow software^52^. Structures were visualized using PyMOL (Schrödinger) and 2D ligand-protein interaction diagrams were generated using LigPlot+^54^.

### Dynamic Light Scattering (DLS)

DLS measurements were performed to analyze the size distribution of AMDHD2 in solution. Directly before measurement, 100 µl protein solution was centrifuged for 10 min at 15,000 g to remove any particles from solution and 70 µl of the supernatant was transferred into a UV disposable cuvette (UVette® 220-1600 nm, Eppendorf #952010051). The cuvette was placed in a Wyatt NanoStar DLS machine and the measurement performed with 10 frames with 10 sec/frame. Data were analyzed with the software Dynamics and converted to particle size distribution functions. The scattering intensity (%) was plotted against the particle radius (nm) in a histogram.

### Baculovirus generation and insect cell expression of GFAT

*Sf21* (DSMZ no. ACC 119) suspension cultures were maintained in SFM4Insect™ HyClone™ medium with glutamine (GE Lifesciences) in shaker flasks at 27°C and 90 rpm in an orbital shaker. GFAT1 and GFAT2 were expressed in *Sf21* cells using the MultiBac baculovirus expression system^55^. In brief, GFAT (from the pFL vector) was integrated into the baculovirus genome via Tn7 transposition and maintained as bacterial artificial chromosome in DH10EMBacY *E. coli* cells. Recombinant baculoviruses were generated by transfection of *Sf21* with bacmid DNA. The obtained baculoviruses were used to induce protein expression in *Sf21* cells.

### GFAT1 and GFAT2 purification

*Sf21* cells were lysed by sonication in lysis buffer (50 mM Tris/HCl pH 7.5, 200 mM NaCl, 10 mM Imidazole, 2 mM TCEP, 0.5 mM Na2Frc6P, 10% (v/v) glycerol) supplemented with complete EDTA-free protease inhibitor cocktail (Roche) and 10 µg/ml DNAseI (Sigma-Aldrich). Cell debris and protein aggregates were removed by centrifugation and the supernatant was loaded on a Ni-NTA Superflow affinity resin (Qiagen). The resin was washed with lysis buffer and the protein eluted with lysis buffer containing 200 mM imidazole. The proteins were further purified according to their size on a HiLoad™ 16/60 Superdex™ 200 prep grade prepacked column (GE Healthcare) using an ÄKTAprime chromatography system at 4°C with a SEC buffer containing 50 mM Tris/HCl, pH 7.5, 2 mM TCEP, 0.5 mM Na2Frc6P, and 10% (v/v) glycerol.

### GDH-coupled activity assay and UDP-GlcNAc inhibition

GFAT’s amidohydrolysis activity was measured with a coupled enzymatic assay using bovine glutamate dehydrogenase (GDH, Sigma-Aldrich G2626) in 96 well standard microplates (F-bottom, BRAND #781602) as previously described^37^ with small modifications. In brief, the reaction mixtures contained 6 mM Frc6P, 1 mM APAD, 1 mM EDTA, 50 mM KCl, 100 mM potassium-phosphate buffer pH 7.5, 6.5 U GDH per 96 well and for L-Gln kinetics varying concentrations of L-Gln. For UDP-GlcNAc inhibition assays the L-Gln concentration was kept at 10 mM. The plate was pre-warmed at 37°C for 10 min and the activity after enzyme addition was monitored continuously at 363 nm in a microplate reader. The amount of formed APADH was calculated with ε_(363 nm, APADH)_ = 9100 l*mol^-1^*cm^-1^. Reaction rates were determined by Excel (Microsoft) and K_m_, v_max_, and IC_50_ were obtained from Michaelis Menten or dose response curves, which were fitted by Prism 8 software (Graphpad).

### GNA1 expression and purification

The expression plasmid for human GNA1 with N-terminal His_6_-tag was cloned previously^15^. Human GNA1 with N-terminal His_6_-tag was expressed in Rosetta (DE3) *E. coli* cells. LB cultures were incubated at 37°C and 180 rpm until an OD_600_ of 0.4-0.6 was reached. Then, protein expression was induced by addition of 0.5 mM IPTG and incubated for 3 h at 37°C and 180 rpm. Cultures were harvested and pellets stored at -80°C. Human GNA1 purification protocol was adopted from Hurtado-Guerrero et al.^56^ with small modifications. *E. coli* were lysed in 50 mM HEPES/NaOH pH 7.2, 500 mM NaCl, 10 mM imidazole, 2 mM 2- mercaptoethanol, 5% (v/v) glycerol with complete EDTA-free protease inhibitor cocktail (Roche) and 10 µg/ml DNAseI (Sigma-Aldrich) by sonication. The lysate was clarified by centrifugation and the supernatant loaded on Ni-NTA Superflow affinity resin (Qiagen). The resin was washed with wash buffer (50 mM HEPES/NaOH pH 7.2, 500 mM NaCl, 50 mM imidazole, 5% (v/v) glycerol) and the protein was eluted with wash buffer containing 250 mM imidazole. Eluted protein was then dialyzed against storage buffer (20 mM HEPES/NaOH pH 7.2, 500 mM NaCl, 5% (v/v) glycerol).

### GNA1 and GNA1-coupled activity assays

The activity of human GNA1 was measured in 96 well standard microplates (F-bottom, BRAND #781602) as described previously^57^. For kinetic measurements, the assay mixture contained 0.5 mM Ac-CoA, 0.5 mM DTNB, 1 mM EDTA, 50 mM Tris/HCl pH 7.5 and varying concentrations of D-GlcN6P. The plates were pre-warmed at 37°C and reactions were initiated by addition of GNA1. The absorbance at 412 nm was followed continuously at 37°C in a microplate reader. The amount of produced TNB, which matches CoA production, was calculated with ε_(412 nm, TNB)_ = 13800 l*mol^-1^*cm^-1^. Typically, GNA1 preparations showed a K_m_ of 0.2 ± 0.1 mM and a kcat of 41 ± 8 sec^-1^.

GFAT’s D-GlcN6P production was measured in a GNA1-coupled activity assay following the consumption of AcCoA at 230 nm in UV transparent 96 well microplates (F-bottom, Brand #781614) as described by Li et al.^57^. In brief, the assay mixture contained 10 mM L-Gln, 0.1 mM AcCoA, 50 mM Tris/HCl pH 7.5, 2 µg hGNA1 and varying concentrations of Frc6P. The plates were incubated at 37°C for 4 min and reactions started by adding L-Gln. Activity was monitored continuously at 230 nm and 37°C in a microplate reader. The amount of consumed AcCoA was calculated with ε_(230 nm, AcCoA)_ = 6436 l*mol^-1^*cm^-1^. As UDP-GlcNAc absorbs light at 230 nm, the GNA-1-coupled assay cannot be used to analyze UDP-GlcNAc effects on activity.

### CRISPR/Cas9-mediated generation of transgenic mice

CRIPSR/Cas9-mediated generation of AMDHD2 knockout mice was performed by ribonucleoprotein complex injection in mouse zygotes. Guide RNAs (crRNAs) targeting exon 4 of the *Amdhd2* locus were designed online (crispor.org) and purchased from IDT. crRNA and tracrRNA were resuspended in injection buffer (1 mM Tris-HCl pH 7.5, 0.1 mM EDTA) and annealed at 1:1 molar concentration in a thermocycler (95°C for 5 min, ramp down to 25°C at 5°C/min). To prepare the injection mix (100 µl), two guide RNAs and the Cas9 enzyme (*S. pyogenes*, NEB) were diluted to a final concentration of 20 ng/µl each in injection buffer. The mix was incubated for 10-15 min at room temperature to allow ribonucleoprotein complex assembly. After centrifugation, 80 µl of the supernatant were passed through a filter (Millipore, UFC30VV25). Both centrifugation steps were performed for 5 min at 13.000 rpm at room temperature. The filtered injection mix was used for zygote injections.

### Mouse Zygote Microinjections

3- to 4-week-old C57Bl/6J females were superovulated by intraperitoneal injection of Pregnant Mare Serum Gonadotropin (5 IU) followed by intraperitoneal injection of Human Chorionic Gonadotropin hormone (5 IU Intervet Germany) 48h later. Superovulated females were mated with 10 to 20 week old stud males. The mated females were euthanized the next day and zygotes were collected in M2 media (Sigma-Aldrich) supplemented with hyaluronidase (Sigma-Aldrich). Fertilized oocytes were injected into the pronuclei or cytoplasma with the prepared CRIPSR/Cas9 reagents. Injections were performed under an inverted microscope (Zeiss AxioObserver) associated micromanipulator (Eppendorf NK2) and the microinjection apparatus (Eppendorf Femtojet) with in-house pulled glass capillaries. Injected zygotes were incubated at 37°C, 5% CO_2_ in KSOM (Merck) until transplantation. 25 zygotes were surgically transferred into one oviduct of pseudo-pregnant CD1 female mice.

All procedures have been performed in our specialized facility, followed all relevant animal welfare guidelines and regulations, and were approved by LANUV NRW 84-02.04.2015.A025.

### Isolation of mouse genomic DNA from ear clips

Ear clips were taken by the Comparative Biology Facility at the Max Planck Institute for Biology of Ageing (Cologne, Germany) at weaning age (3-4 weeks of age) and stored at -20°C until use. 150 µl ddH2O and 150 µl directPCR Tail Lysis reagent (Peqlab) were mixed with 3 µl proteinase K (20 mg/ml in 25 mM Tris-HCl, 5 mM Ca_2_Cl, pH 8.0, Sigma-Aldrich). This mixture was applied to the ear clips, which were then incubated at 56°C overnight (maximum 16 h) shaking at 300 rpm. Proteinase K was inactivated at 85°C for 45 min without shaking. The lysis reaction (2 µl) was used for genotyping PCR without further processing. For genotyping of mouse genomic DNA DreamTaq DNA polymerase (ThermoFisher Scientific) was used.

### Alignments

Following UnitProt IDs were used for the protein sequence alignment of AMDHD2: *Homo sapiens* isoform 1: Q9Y303-1, *Mus musculus*: Q8JZV7, *Caenorhabditis elegans*: P34480, *Candida albicans*: Q9C0N5, *Escherichia coli*: P0AF18, *Bacillus subtilis*: O34450. ClustalOmega (https://www.ebi.ac.uk/Tools/msa/clustalo/) was used to generate a multiple sequence alignment^58^. The alignment was formatted with the ESPript3 server (*espript.ibcp.fr/)*^59^ and further modified.

### Statistical analysis

Data are presented as mean ± SEM or as mean + SEM. The mean of technical replicates is plotted for each biological replicate. Biological replicates represent different passages of the cells that were seeded on independent days. Statistical significance was calculated using GraphPad Prism (GraphPad Software, San Diego, California). The statistical test used is indicated in the respective figure legend. Significance levels are * p<0.05, ** p<0.01, *** p<0.001 versus the respective control.

### Data availability

Structural data reported in this study have been deposited in the Protein Data Bank with the accession codes 7NUT [https://doi.org/10.2210/pdb7NUT/pdb] and 7NUU [https://doi.org/10.2210/pdb7NUU/pdb]. All other data supporting the presented findings are available from the corresponding authors upon request.

## Acknowledgments

We thank all M.S.D. and U.B. laboratory members as well as L. Kurian, H. Bazzi, and D. Vilchez for helpful discussions. The FLAG-HA-hGFAT-2-pcDNA3.1 plasmid was kindly provided by C. Geisen (Max Planck Institute for Biology of Ageing, MPI-AGE). We thank Y. Hinze, S. Perin and P. Giavalisco from the MPI-AGE metabolomics core facility. We thank K. Folz-Donahue, L. Schumacher, A. Just, and C. Kukat from the MPI-AGE FACS and imaging core facility. We thank F. Metge, and J. Boucas from the MPI-AGE bioinformatics core facility. We thank I. Vogt from the MPI-AGE transgenesis core facility. We thank the MPI-AGE comparative biology facility. We thank M. Grzonka for support with the E14 mESCs. We are grateful to S. Birkmann for support in the insect cell maintenance. We thank I. Grimm and S. Schäfer for their help with the AMDHD2 production. Crystals were grown in the Cologne Crystallization facility (C_2_f). We thank the staff of beamline X06SA at the Swiss Light Source, Paul Scherrer Institute, Villigen (Switzerland) and beamline P13 at PETRA III, DESY, Hamburg (Germany) for their support during data collection. This work was supported by the German Federal Ministry of Education and Research (BMBF, grant 01GQ1423A EndoProtect), by the German Research Foundation (DFG, Projektnummer 73111208-SFB 829, B11 and B14), by the European Commission (ERC-2014-StG-640254-MetAGEn), and by the Max Planck Society. The Cologne Crystallization Facility C_2_f was supported by DFG grant INST 216/949-1 FUGG.

## Author contributions

V.K, S.R., and M.S.D. designed the project. M.S.D. and U.B. supervised the project. V.K., K.A., S.M., and M.H. performed the genetic and cell biology work in Figures 1 and 4. S.R. performed the biochemical and crystallization experiments in Figures 2 and 3. S.R. and S.B. did the activity assays in Figure 4. L.E. and B.S. generated the AMDHD2 antibody. V.K, S.R., and M.S.D. wrote the manuscript. V.K. and S.R. prepared the figures.

## List of Figure supplements

**Figure 1-figure supplements 1 and 2.**

**Figure 2-figure supplements 1-8**

**Figure 4-figure supplements 1-4**

## List of source data

**Figure 1- source data 2:** anti-AMDHD2 Western Blot (labelled) (Figure 1C)

**Figure 1- source data 3:** anti-Tubulin Western Blot (raw) (Figure 1C)

**Figure 1- source data 4:** anti-Tubulin Western Blot (labelled) (Figure 1C)

**Figure 1- source data 5:** XTT assay of WT and AMDHD2 K.O. AN3-12 cells (Figure 1D)

**Figure 1- source data 6:** IC-MS analysis of UDP-GlcNAc levels (Figure 1G)

**Figure 1-figure supplement 1-source data 1:** XTT assay of WT and AMDHD2 K.O. AN3-12 cells (Figure 1- figure supplement 1A)

**Figure 1-figure supplement 1-source data 2:** XTT assay of WT and AMDHD2 K.O. AN3-12 cells (Figure 1- figure supplement 1C)

**Figure 1-figure supplement 1-source data 3:** IC-MS analysis of UDP-GlcNAc levels (Figure 1- figure supplement 1D)

**Figure 2-source data 1:** Thermal Shift Assay, melting temperatures of AMDHD2 in SEC buffer in the presence of varying divalents (Figure 2a)

**Figure 2-source data 2:** Deacetylase activity of AMDHD2 in the presence of EDTA and several indicated divalents (Figure 2b)

**Figure 2-figure supplement 3-source data 1**: Representative size-exclusion chromatography of AMDHD2 (Figure 2-figure supplement 3A)

**Figure 2-figure supplement 3-source data 2:** Representative dynamic light scattering measurement of AMDHD2 (Figure 2-figure supplement 3B)

**Figure 3-source data 1-4:** SDS-gels stained with Coomassie brilliant blue of a representative bacterial test expression of the human AMDHD2 variants in an Excel file (full raw unedited and labeled) (Figure 3a)

**Figure 3-source data 1:** First SDS-gel stained with Coomassie brilliant, provided as full raw unedited and labeled JPG files (Figure 3a)

**Figure 3-source data 2:** Second SDS-gel stained with Coomassie brilliant, provided as full raw unedited and labeled JPG files (Figure 3a)

**Figure 3-source data 3:** Third SDS-gel stained with Coomassie brilliant, provided as full raw unedited and labeled JPG files (Figure 3a)

**Figure 3-source data 4:**, Fourth SDS-gel stained with Coomassie brilliant, provided as full raw unedited and labeled JPG files (Figure 3a)

**Figure 3-source data 5:** Deacetylase activity of wild type (WT) and mutant human AMDHD2 (Figure 3c)

**Figure 4- source data 1:** Relative Gfat1 and Gfat2 mRNA levels (qPCR) in WT AN3-12 cells (Figure 4A)

**Figure 4- source data 2:** anti-GFAT1 Western Blot (raw) (Figure 4B)

**Figure 4- source data 3:** anti-GFAT1 Western Blot (labelled) (Figure 4B)

**Figure 4- source data 4:** anti-Tubulin Western Blot (raw) (Figure 4B)

**Figure 4- source data 5:** anti-Tubulin Western Blot (labelled) (Figure 4B)

**Figure 4- source data 6:** anti-GFAT2 Western Blot (raw) (Figure 4B)

**Figure 4- source data 7:** anti-GFAT2 Western Blot (labelled) (Figure 4B)

**Figure 4- source data 8:** anti-Tubulin Western Blot (raw) (Figure 4B)

**Figure 4- source data 9:** anti-Tubulin Western Blot (labelled) (Figure 4B)

**Figure 4- source data 10:** anti-AMDHD2 Western Blot (raw) (Figure 4C)

**Figure 4- source data 11:** anti-AMDHD2 Western Blot (labelled) (Figure 4C)

**Anti-TUBULIN:** same as Figure 4- source data 4 (Figure 4C)

**Anti-TUBULIN:** same as Figure 4- source data 5 (Figure 4C)

**Figure 4- source data 12:** IC-MS analysis of UDP-GlcNAc levels of AMDHD2 K.O. compared to WT cells (Figure 4D)

**Figure 4- source data 13:** UDP-GlcNAc dose-response assay with hGFAT1 and hGFAT2 (Figure 4E)

**Figure 4- source data 14:** Relative UDP-GlcNAc levels in indicated cell lines measured by IC-MS (Figure 4F)

**Figure 4- source data 15:** anti-GFAT2 Western Blot (raw) (Figure 4G)

**Figure 4- source data 16:** anti-GFAT2 Western Blot (labelled) (Figure 4G)

**Figure 4- source data 17:** anti-Tubulin Western Blot (raw) (Figure 4G)

**Figure 4- source data 18:** anti-Tubulin Western Blot (labelled) (Figure 4G)

**Figure 4- source data 19:** Relative Gfat2 mRNA level (qPCR) in WT AN3-12 cells and upon partial differentiation by a 5-day LIF removal (-Lif) (Figure 4H)

**Figure 4- figure supplement 1-source data 1:** Relative UDP-GlcNAc levels of distinct mutants compared to WT AN3-12 mESCs (Figure 1- figure supplement1)

**Figure 4-figure supplement 2-source data 1:** L-Gln kinetic of WT human GFAT1 and WT human GFAT2 (Figure 4- figure supplement 2A)

**Figure 4-figure supplement 2-source data 2:** Fructose-6-Phosphate kinetic of WT human GFAT1 and WT human GFAT2 (Figure 4- figure supplement 2B)

**Figure 4-figure supplement 3-source data 1:** anti-RL2 Western Blot (raw) (Figure 4- figure supplement 3)

**Figure 4- figure supplement 3-source data 2:** anti-GFAT2 Western Blot (labelled) (Figure 4- figure supplement 3)

**Anti-TUBULIN:** same as Figure 4- source data 8 (Figure 4- figure supplement 3)

**Anti-TUBULIN:** same as Figure 4- source data 9 (Figure 4- figure supplement 3)

**Figure 4- figure supplement 4-source data 1**: Relative Nanog and Klf4 mRNA level (qPCR) in WT AN3-12 cells and upon partial differentiation by a 5-day LIF removal (-Lif) (Figure 4-Fig. Supp. 4A)

**Figure 4- figure supplement 4-source data 2**: Relative Gfat1 and Amdhd2 mRNA level (qPCR) in WT AN3-12 cells and upon partial differentiation by a 5- day LIF removal (-Lif) (Figure 4-Fig. Supp. 4B)

**Figure 4- figure supplement 4-source data 3**: anti-GFAT1 Western Blot (raw) (Fig. 4-Fig Supp. 4C)

**Figure 4- figure supplement 4-source data 4**: anti-GFAT1 Western Blot (labelled) (Fig. 4-Fig Supp. 4C)

**Figure 4- figure supplement 4-source data 5**: anti-AMDHD2 Western Blot (raw) (Fig. 4-Fig Supp. 4C)

**Figure 4- figure supplement 4-source data 6**: anti-AMDHD2 Western Blot (labelled) (Fig. 4-Fig Supp. 4C)

**Figure 4- figure supplement 4-source data 7**: anti-TUBULIN Western Blot (raw) (Fig. 4-Fig Supp. 4C)

**Figure 4- figure supplement 4-source data 8**: anti-TUBULIN Western Blot (labelled) (Fig. 4-Fig Supp. 4C)

**Figure 1- source data 1:** anti-AMDHD2 Western Blot (raw) (Figure 1C)

---

**Figure 1-figure supplement 1.**
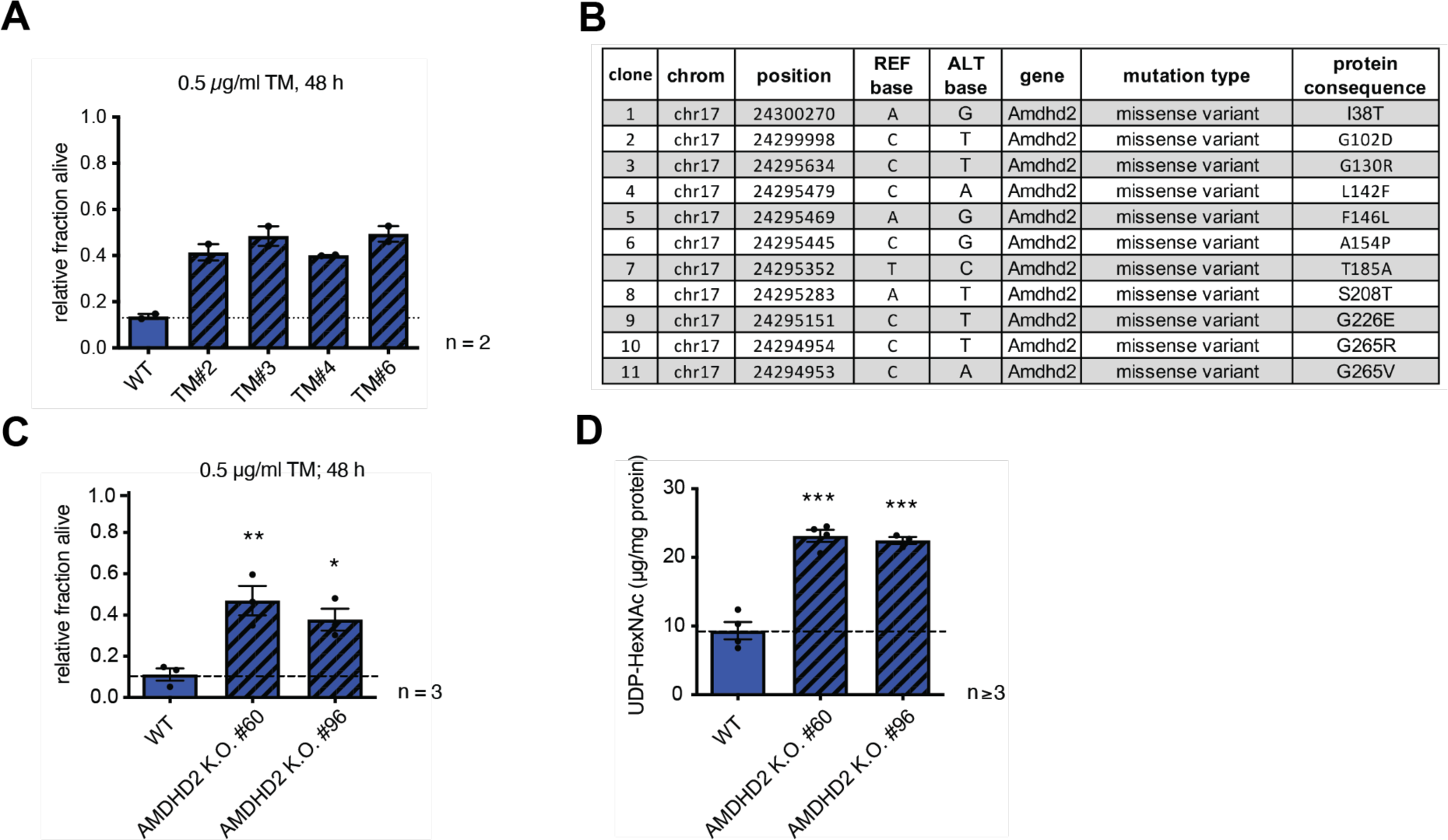
Chemical mutagenesis screen for tunicamycin resistance in mESCs identifies AMDHD2. Identification and confirmation of AMDHD2 as the causative gene for mediating TM resistance by elevated UDP-GlcNAc levels. **(A)** Cell viability (XTT assay) of four TM resistant AN3-12 clones from mutagenesis screen that were used for WES upon treatment with 0.5 µg/ml TM for 48h (mean ± SEM, n=2). **(B)** Table listing all mutations in the *Amdhd2* locus identified in the TM resistance screen. **(C)** Cell viability (XTT assay) of WT and two additional independently generated AMDHD2 K.O. AN3-12 cell lines upon treatment with 0.5 µg/ml TM for 48h (mean ± SEM, n=3, * p<0.05; ** p<0.01, one-way ANOVA). **(D)** UDP-HexNAc concentration of the two additional AMDHD2 K.O. cell lines compared to WT AN3-12 mESCs (mean ± SEM, n≥3, *** p<0.001, one-way ANOVA). UDP-HexNAc is the combined pool of the UDP-GlcNAc and UDP-GalNAc epimer pools.

**Figure 1-figure supplement 2.**
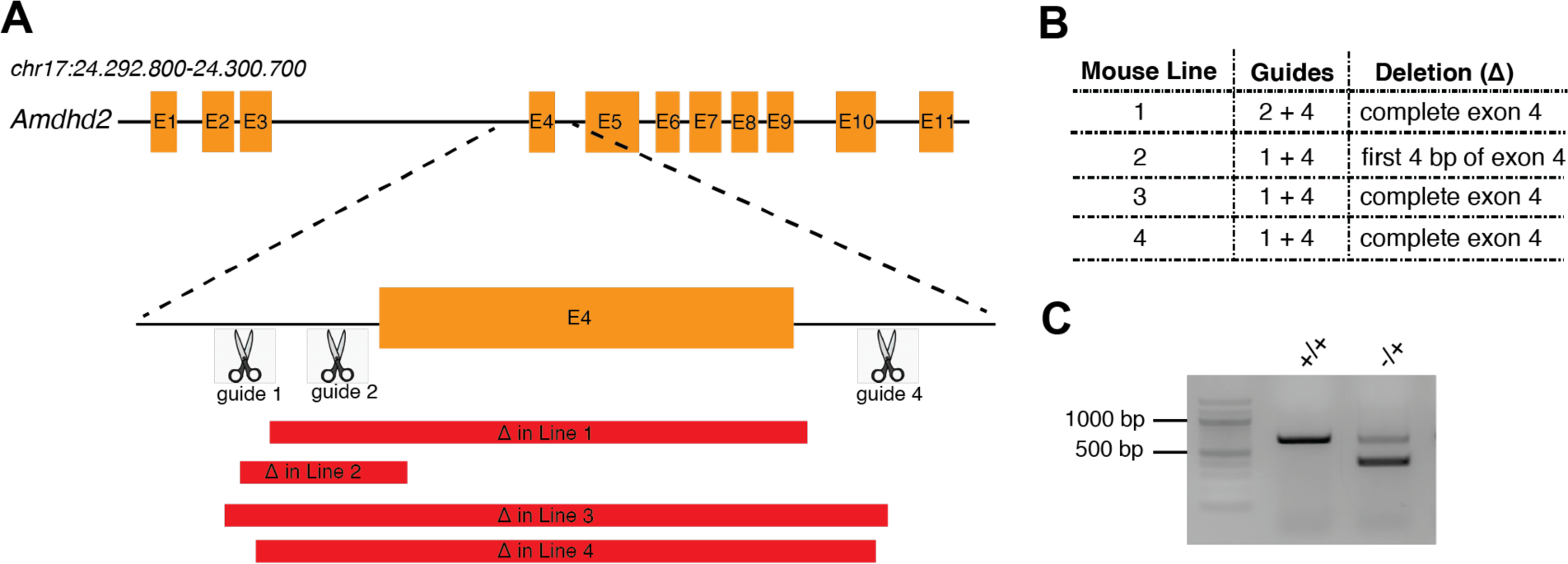
Structural and biochemical characterization of human AMDHD2. Generation of different AMDHD2 K.O. founder mice. **(A)** Schematic of the CRISPR/Cas9 targeted exon of the mouse *Amdhd2* locus. Deletions in founder lines 1-4 are indicated in red. **(B)** Table listing used guide combinations and deletion details of the AMDHD2 K.O. founder lines. **(C)** Representative genotyping results of AMDHD2 K.O. mice. The WT PCR product is 675 bp and the *Amdhd2* K.O. allele shows a size of 300 bp (line 902).

**Figure 2-figure supplement 1.**
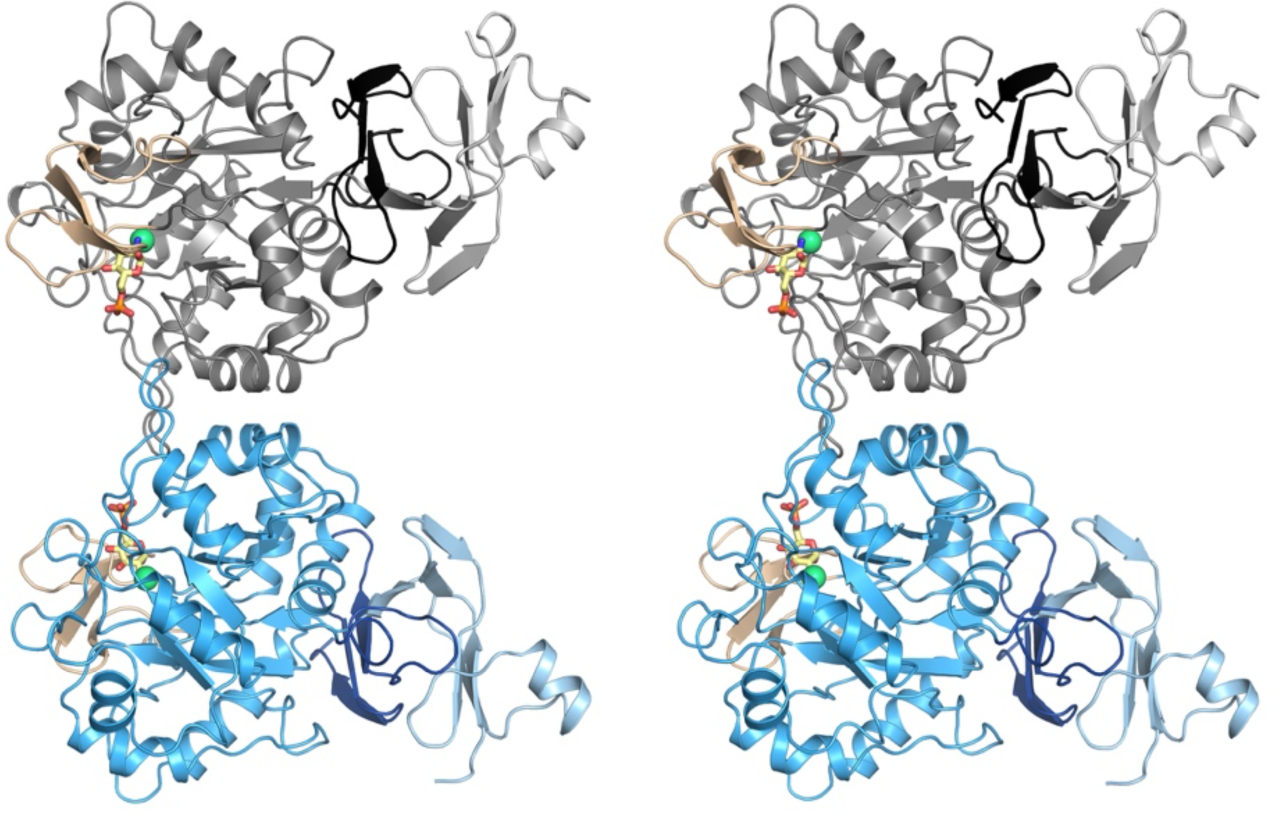
Structural and biochemical characterization of human AMDHD2. Stereo image of the human AMDHD2 dimer in cartoon representation. Monomer A is colored in gray and monomer B in blue. The two deacetylase domains are interacting with each other. The DUF domain is formed by residues of the N-terminus (light gray, light blue) and residues of the C-terminus (black, dark blue). GlcN6P (yellow sticks), Zn^2+^ (green sphere), and the putative active site lid (wheat) are highlighted.

**Figure 2-figure supplement 2.**
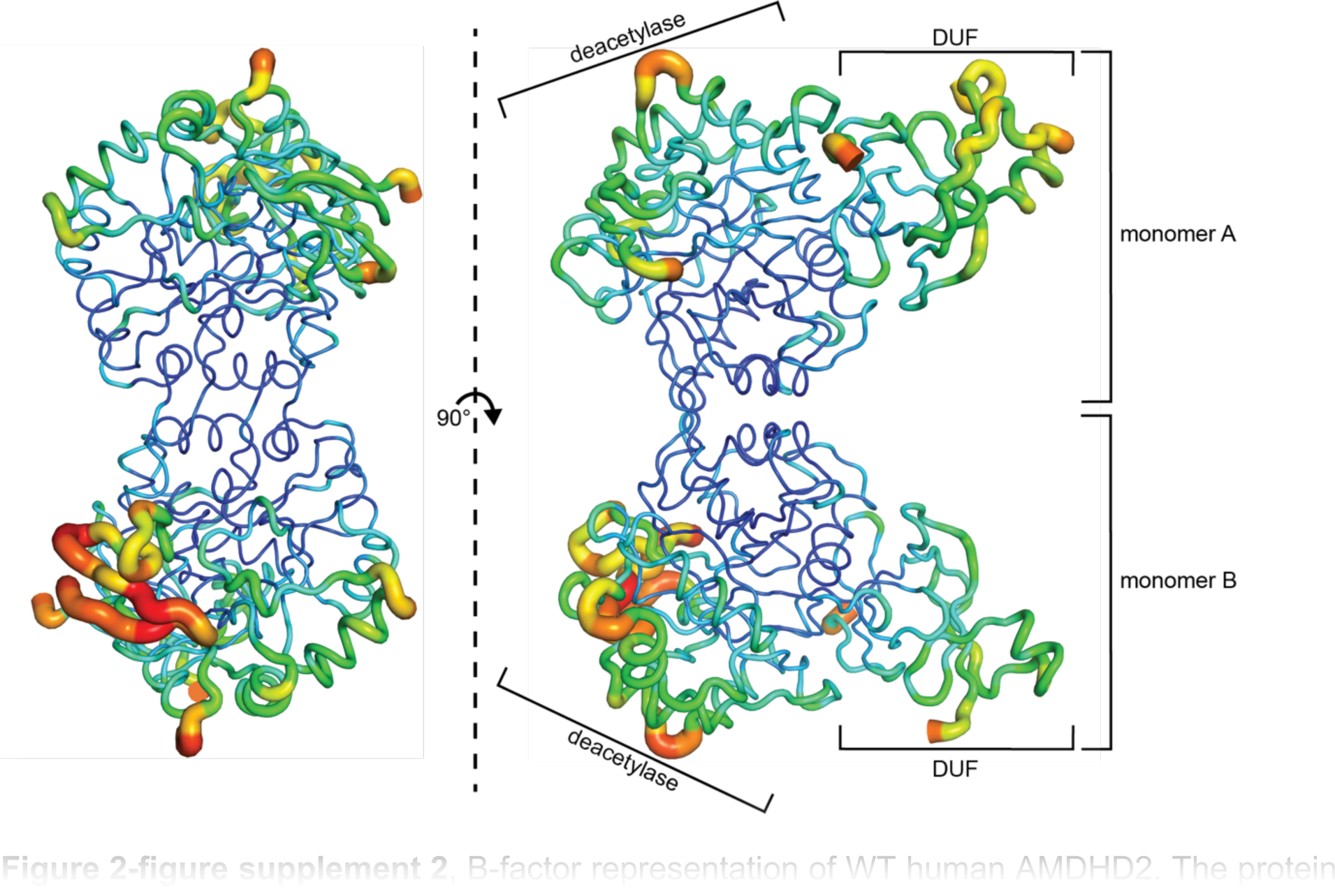
B-factor representation of WT human AMDHD2. The protein is presented as putty cartoon and colored from low to high B-factors (24-108 A^2^, blue to red).

**Figure 2-figure supplement 3.**
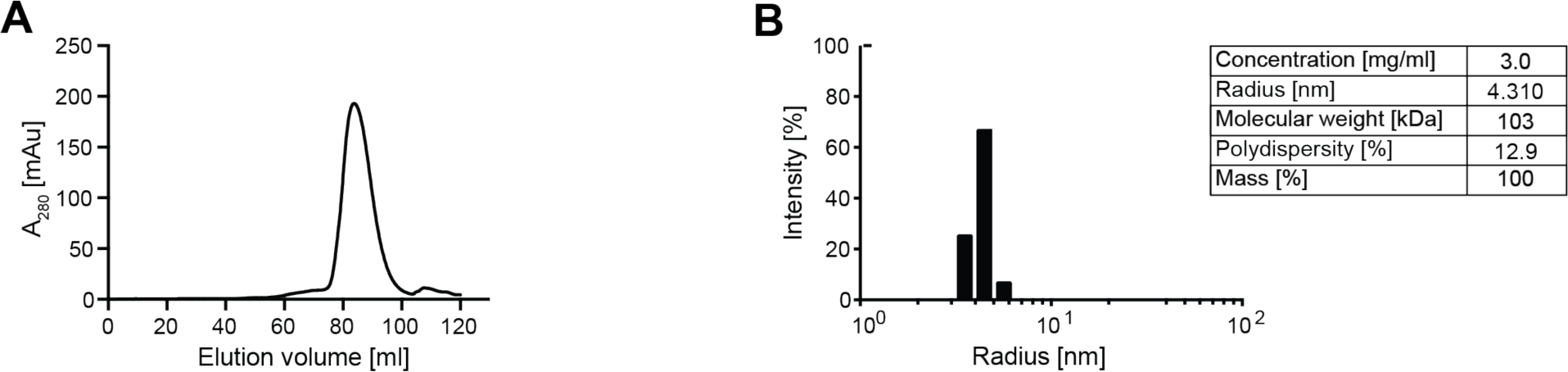
Oligomeric state of human AMDHD2. (A) Representative chromatogram of a size exclusion chromatography of human AMDHD2 using a HiLoad Superdex 200 16/600 column. Absorption at 280 nm (mAU: milli absorbance units) was plotted against the elution volume. AMDHD2 elutes as monomer. **(B**) Representative DLS measurement of WT AMDHD2. Table: Parameters of the representative DLS measurement showing a dimeric assembly.

**Figure 2-figure supplement 4.**
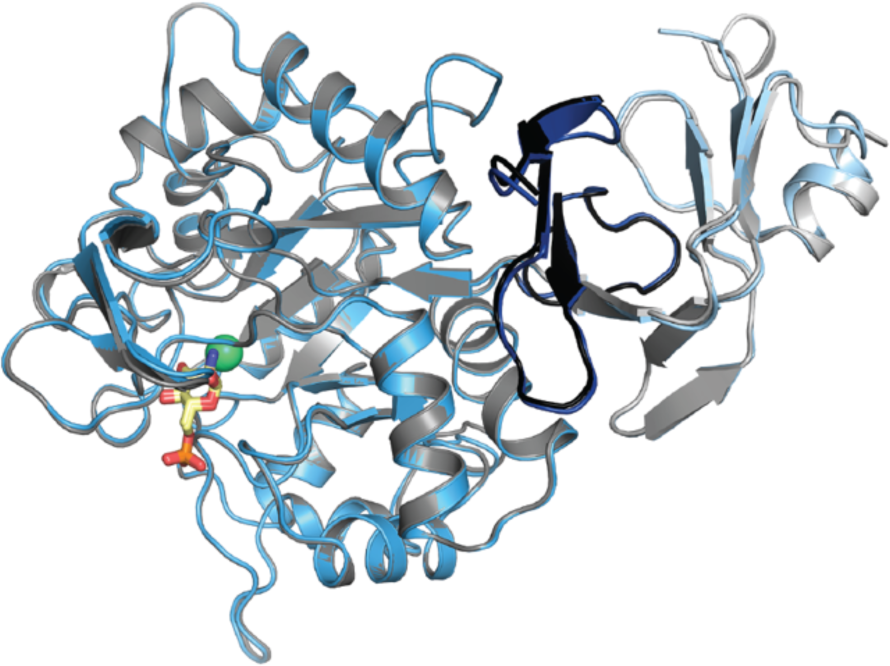
Superposition of GlcN6P-bound AMDHD2 monomer A (gray) and monomer B (blue) in cartoon representation. GlcN6P (yellow sticks) and Zn^2+^ (green spheres) are highlighted.

**Figure 2-figure supplement 5.**
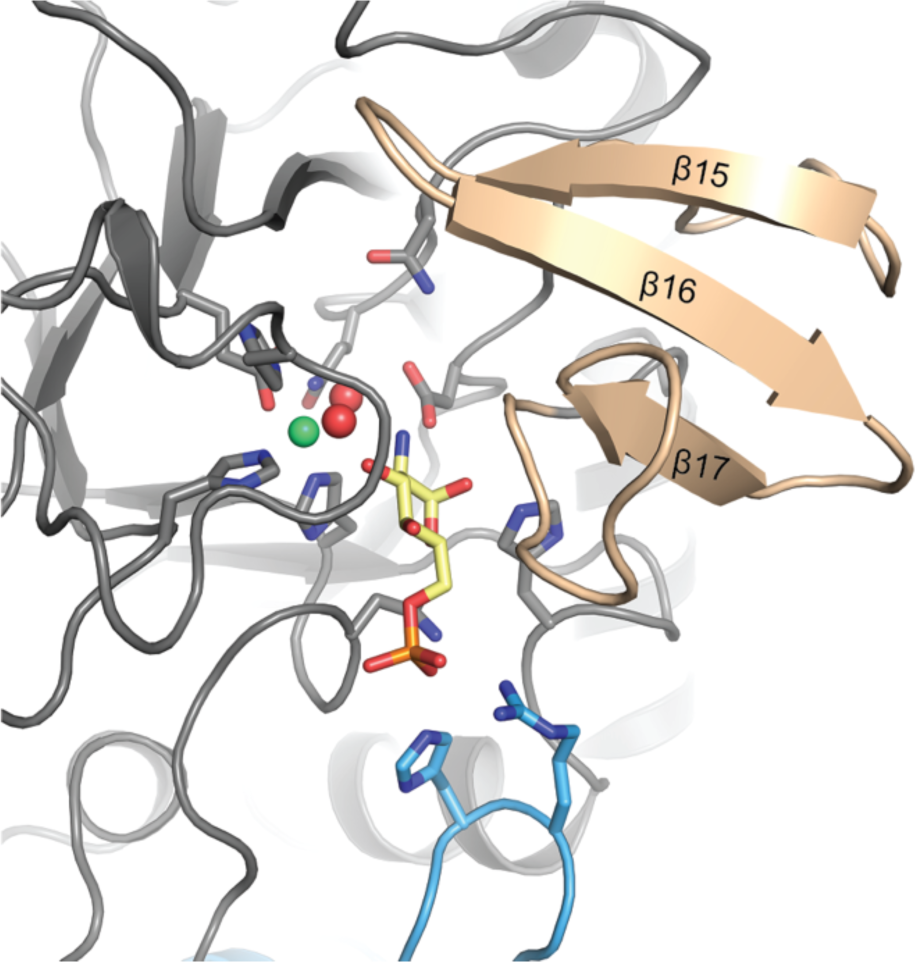
Close-up view of the active site in cartoon representation. Residues involved in ligand binding or catalysis are highlighted as sticks, as well as GlcN6P (yellow sticks), Zn^2+^ (green sphere) and two water molecules (red spheres). Three antiparallel β-strands (β15-β17, colored wheat) build a β-sheet close to the active site.

**Figure 2-figure supplement 6.**
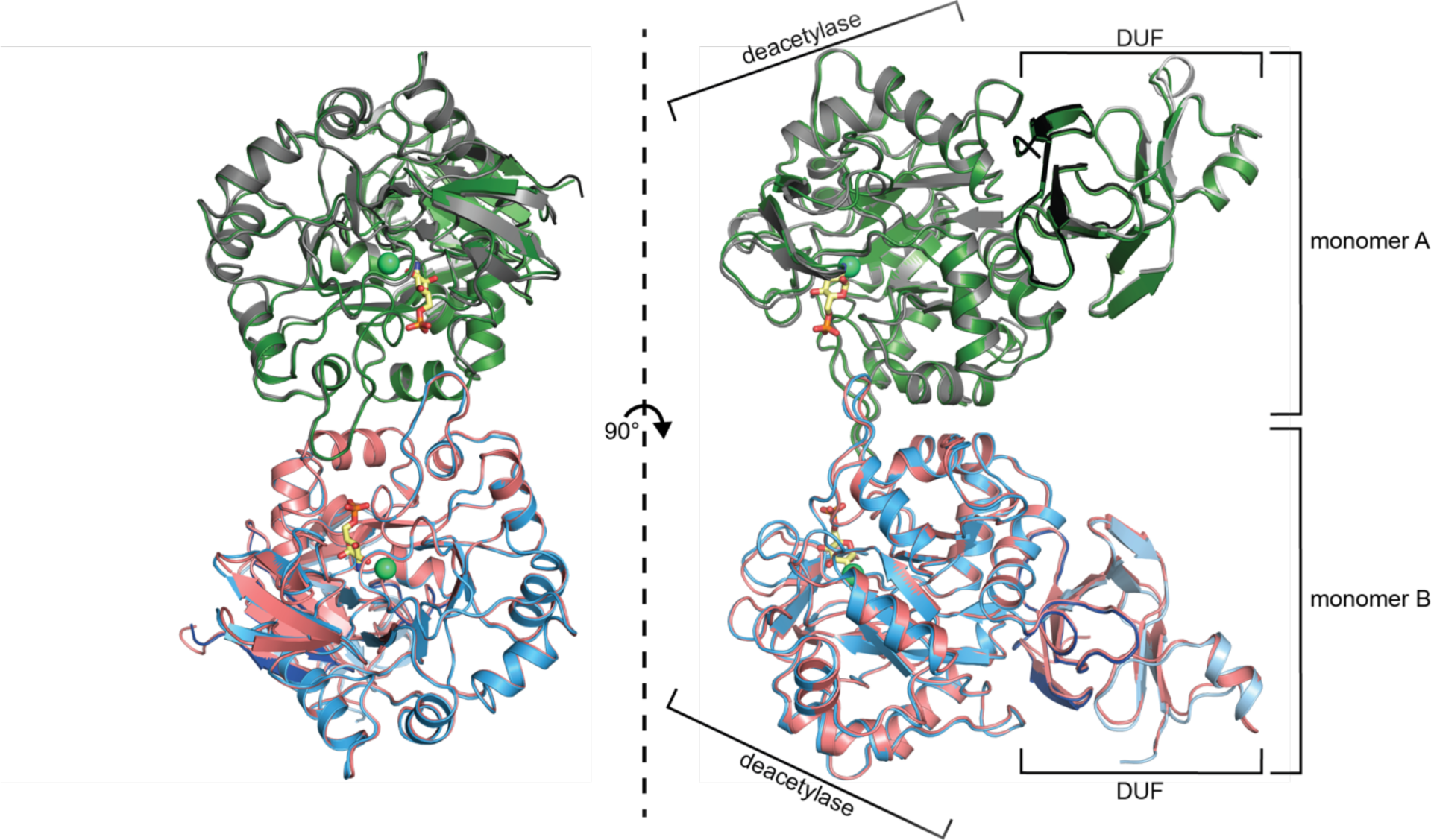
Superposition of the structures of GlcN6P-bound (gray, blue) and GlcN6P-free (green, red) human AMDHD2 with RMSD of 0.67 Å over 792 main chain residues in cartoon representation. GlcN6P (yellow sticks) and Zn^2+^ (green spheres) are highlighted.

**Figure 2-figure supplement 7.**
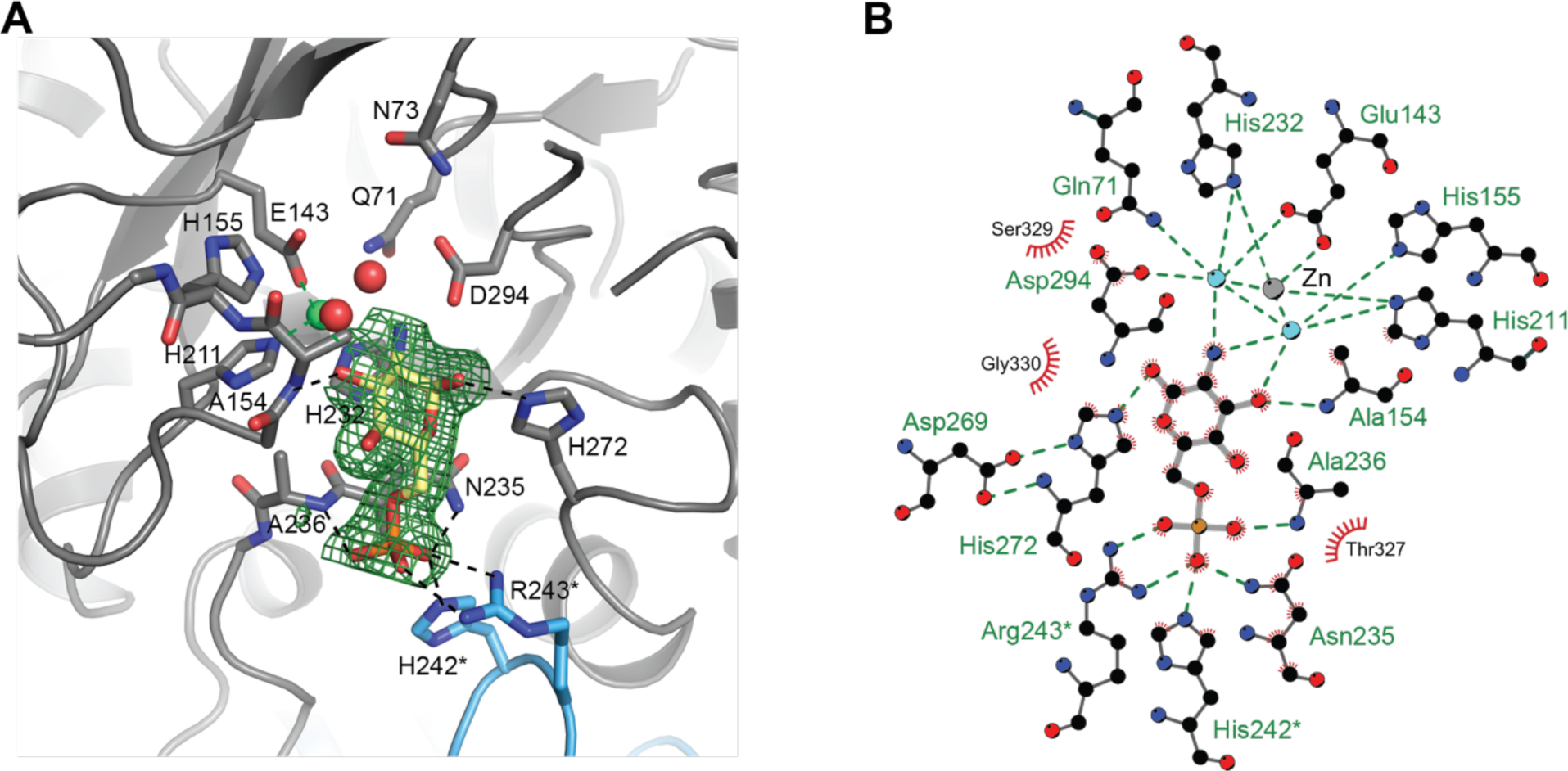
Active site of human AMDHD2. (**A**) Omit map of the active site of human AMDHD2 in cartoon representation. Residues involved in ligand binding or catalysis are highlighted as sticks, as well as GlcN6P (yellow sticks). The Fo-Fc omit map is colored in green and its contour level is at 1.5 RMSD. (**B**) 2D ligand-protein interaction diagram of GlcN6P interacting with human AMDHD2. Ligand bonds are colored in gray and amino acid side chain bonds in black. Green dashed lines indicate hydrogen bonds and red spiked arcs present residues making non-bonded contacts with the ligands.

**Figure 2-figure supplement 8.**
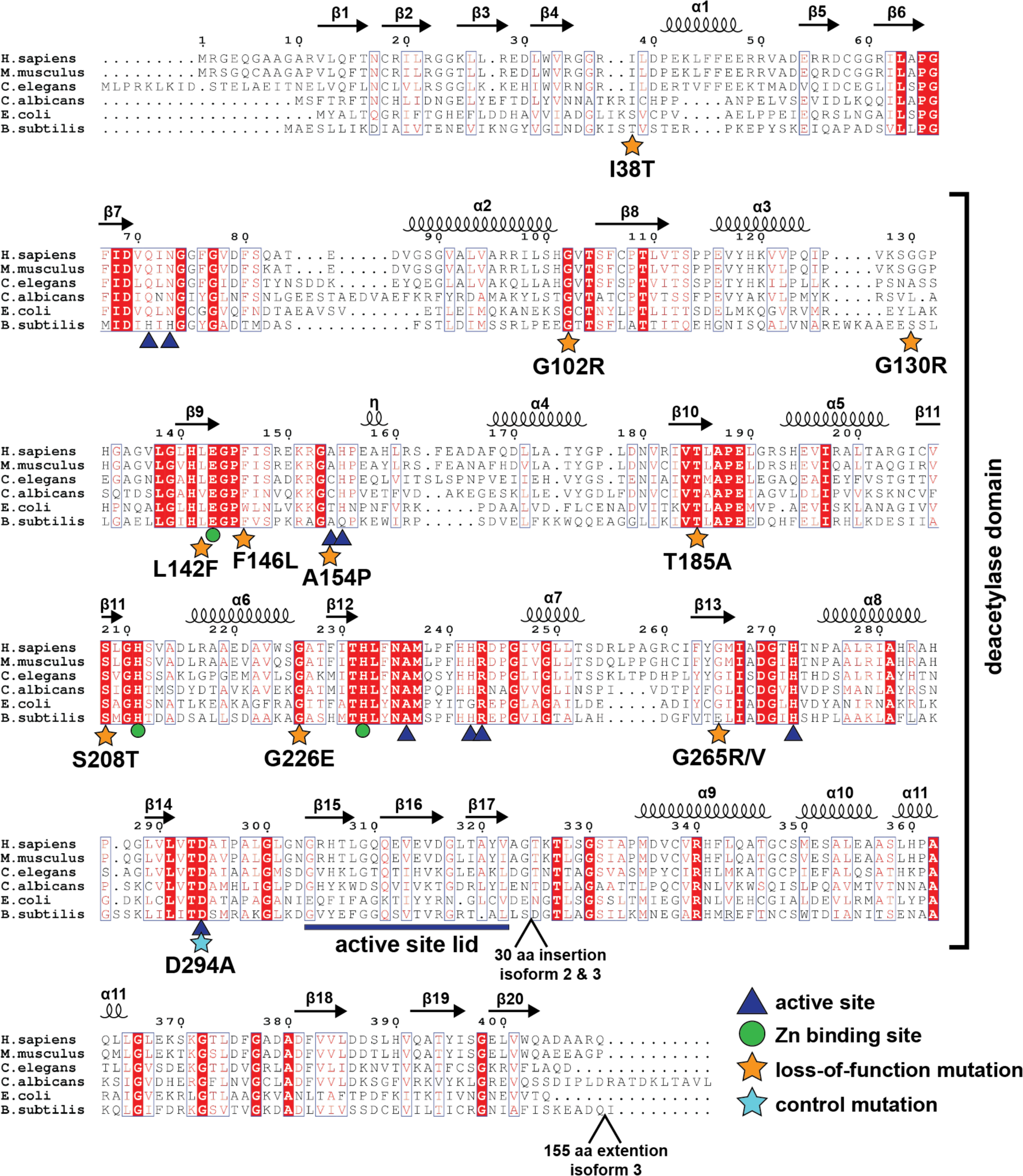
Protein sequence alignment of AMDHD2. Red boxes indicate identical residues, red letters indicate similar residues. The deacetylase domain and secondary structure elements are annotated, as well as the positions of insertions and extensions in AMDHD2 isoform 2 and isoform 3. Residues involved in product binding, catalysis, or metal binding are highlighted. The putative active site lid is marked. Moreover, the positions of the potential loss-of-function mutations and the control mutation are labeled.

**Figure 4-figure supplement 1.**
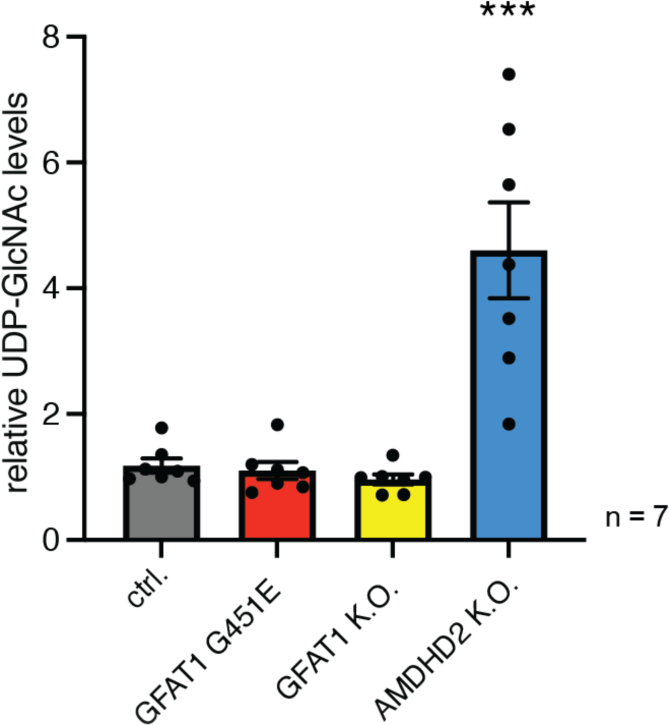
AMDHD2 limits HBP activity when GFAT2 replaces GFAT1 as the first enzyme. Manipulation of GFAT1 in AN3-12 cells has no influence on UDP-GlcNAc levels. IC-MS analysis of relative UDP-GlcNAc levels of WT, GFAT1 G451E, GFAT1 K.O. and AMDHD2 K.O. AN3-12 cells. (mean ± SEM, n=7, *** p<0.001, one-way ANOVA)

**Figure 4-figure supplement 2.**
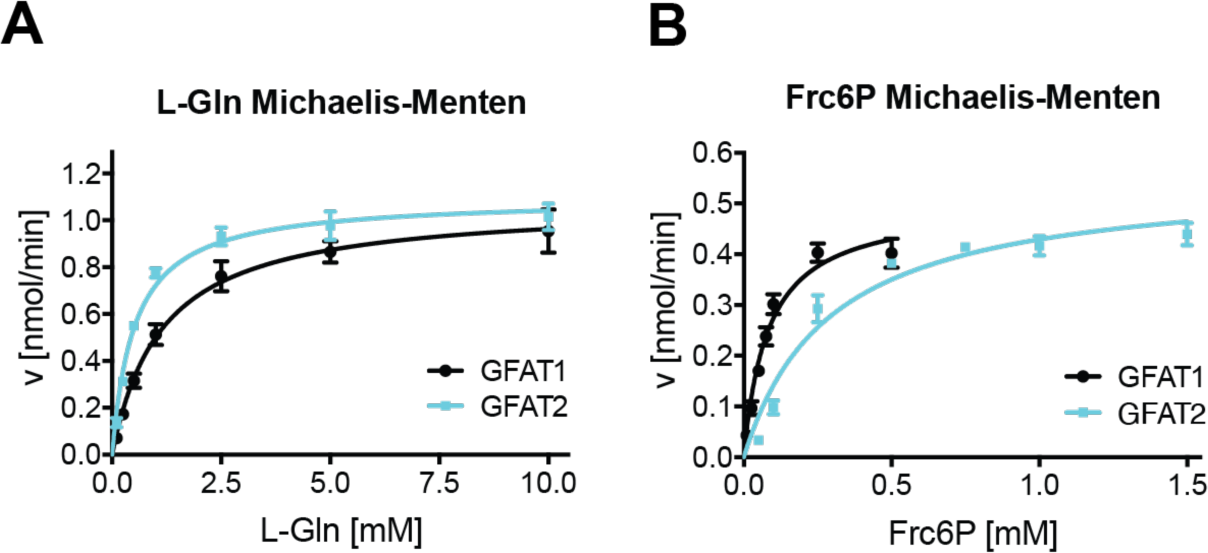
Biochemical characterization of human GFAT2 compared to human GFAT1. **(A)** L-Gln kinetic of WT human GFAT1 (black circle) and WT human GFAT2 (teal square) (mean ± SEM, hGFAT1 n=5, hGFAT2 n=4). **(B)** Frc6P kinetic of WT hGFAT1 (black circle) and WT hGFAT2 (teal square) (mean ± SEM, hGFAT1 n=5, hGFAT2 n=8).

**Figure 4-figure supplement 3.**
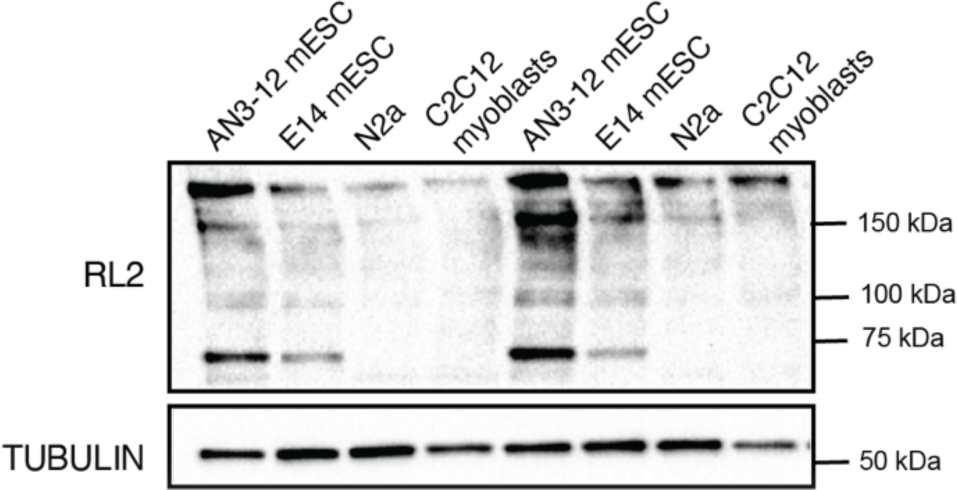
O-GlcNAcylation levels of different cell lines. Western blot analysis of O-GlcNAc-modified proteins (RL2) in the indicated cell lines.

**Figure 4-figure supplement 4.**
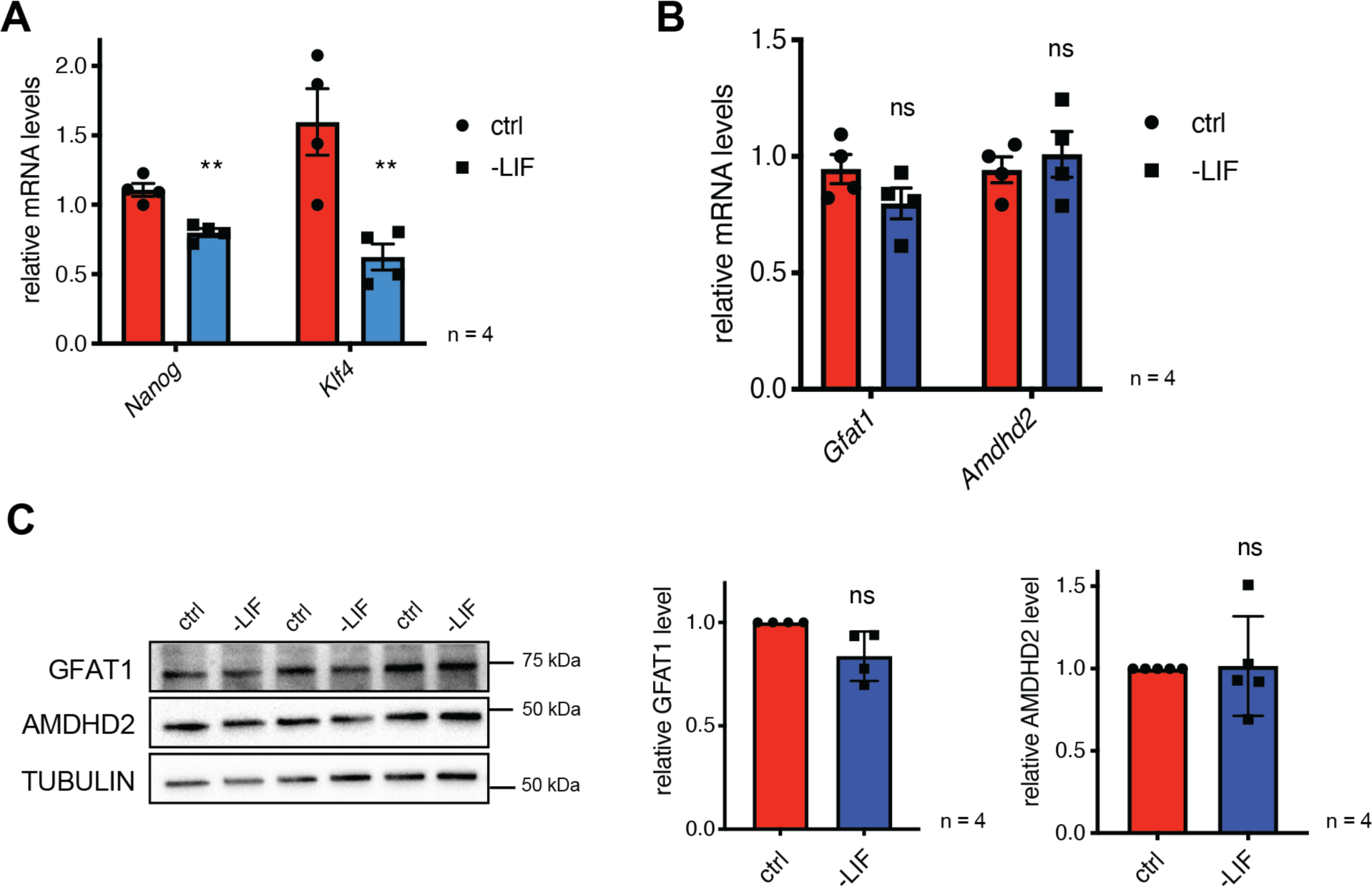
The effect of partial differentiation upon LIF removal in AN3- 12 cells on the enzymatic HBP composition**. (A)** Relative *Nanog* and *Klf4* mRNA-level (qPCR) of WT AN3-12 cells and upon partial differentiation by 5 days of LIF removal (SEM ± n=4, ns = not significant, unpaired t-test). **(B)** Relative *Gfat1* and *Amdhd2* mRNA-level (qPCR) of WT AN3-12 cells and upon partial differentiation by 5 days of LIF removal (mean ± SEM, n=4, ns = not significant, unpaired t-test). **(C)** Western blot analysis of GFAT1 and AMDHD2 in WT AN3-12 cells and upon partial differentiation by 5 days of LIF removal, including quantification relative to tubulin and the WT control cells (mean ± SD, n=4, ns = not significant, unpaired t-test).

